# The U1 snRNP protein U1C and Helix H of U1 snRNA are critical for small molecule splicing modulator function

**DOI:** 10.1101/2025.08.21.671662

**Authors:** Zhiling Kuang, Xueni Li, Zhichao Tang, Brian Kosmyna, Shasha Shi, Karoline Lambert, Wenzheng Zhang, Joseph Giovinazzo, Kerstin A. Effenberger, Jairo Sierra, Lanqing Ying, Scott J. Barraza, Wencheng Li, Christopher R. Trotta, Jingxin Wang, Rui Zhao

**Affiliations:** Department of Biochemistry and Molecular Genetics, University of Colorado Anschutz Medical Campus; Section of Genetic Medicine, Department of Medicine, Biological Sciences Division, University of Chicago; PTC Therapeutics, Inc., Warren, NJ 07059, USA

## Abstract

Risdiplam and branaplam represent two classes of small-molecule splicing modulators that promote U1 snRNP recognition of weak non-canonical GA/GU-containing 5’ splice sites (ss). We demonstrated that branaplam enhanced recognition of these 5’ ss by reconstituted U1 snRNP in vitro, and that this effect depended on the ZnF domain of U1C and Helix H of U1 snRNA, but not U1A or U1-70K. In cells, depletion of U1C generally reduced compound-induced exon inclusion for most cassette exons. Interestingly, a subset of cassette exons became responsive to compound only upon U1C knockdown, supporting a model in which U1C stabilizes specific conformations at the 5’ ss/U1 snRNA interface in a context-dependent manner that can either facilitate or hinder compound binding. Surprisingly, risdiplam shows no effect on weak 5’ ss recognition in vitro, suggesting additional cellular factors are required for its activity.

## Introduction

Pre-mRNA splicing is an essential step in the gene expression pathway of all eukaryotes ^1,2^. A majority of human genes contain alternatively spliced introns, which serve as a fundamental mechanism of gene regulation ^3^. Accurate and efficient recognition of the splice sites is crucial for eukaryotic gene expression. The spliceosome, a huge protein-RNA complex, is responsible for recognizing and splicing introns ^4^. During early spliceosome assembly, the 5’ splice site (ss) is recognized by the U1 snRNP, a major component of the spliceosome ^5–8^. Human U1 snRNP consists of a 164 nt U1 snRNA, seven Sm proteins that form a heptameric Sm ring, U1A, U1C, and U1-70K proteins. The recognition of the 5’ ss by U1 snRNP involves base pairing between the 5’ ss and the 5’ end of U1 snRNA. The U1C protein binds and stabilizes this RNA duplex using its Zinc-Finger (ZnF) domain ^9,10^. Recently, small molecule splicing modulators that improve the recognition of the weak non-canonical 5’ ss by U1 snRNP have emerged as a unique approach to efficiently modulate protein levels in genetic diseases affecting the CNS, such as Spinal Muscular Atrophy (SMA).

SMA is the leading cause of infant death in the US. It is caused by the mutation or deletion of the SMN1 gene resulting in the reduction or absence of SMN proteins and subsequent damage or death of motor neurons ^11^. All humans have a paralog of the gene, SMN2, which encodes the same protein but contains silent mutations that lead to the skipping of exon 7, which contains a weak non-canonical 5’ ss with the <u>GA</u>/GU sequence (where “/” denotes the exon and intron boundary and nucleotides upstream and downstream of the boundary are numbered as -1 and +1, respectively) instead of the canonical <u>AG</u>/GU sequence ^12,13^. The skipping of exon 7 generates a truncated and non-functional protein. Inhibition of this skipping increases the level of full-length SMN protein and rescues disease phenotypes in mouse models ^14^. PTC Therapeutics, in collaboration with Roche and the SMA Foundation, identified and developed the first class of sequence-specific small molecule splicing modulators represented by risdiplam. These modulators selectively improve the recognition of the non-canonical GA/GU 5’ ss by U1 snRNP^14^. Risdiplam is orally bioavailable with excellent CNS distribution and has been approved for the treatment of SMA by the FDA since 2020.

Risdiplam and related compounds have generated tremendous excitement in the field as small-molecule splicing modulators offer new approaches to modulating protein levels. One such approach is to promote the inclusion of an exon with a non-canonical 5’ ss by a small molecule in an intron of a pathogenic protein that creates a premature stop codon, leading to reduced protein levels through nonsense-mediated mRNA decay (NMD) ^15,16^{Fair, 2024 #4347}. This approach is particularly attractive for neurodegenerative diseases such as Huntington’s disease due to the excellent distribution profile of these small molecules in the CNS. Two small-molecule splicing modulators, HTT-C2 and branaplam, target the GA/GU 5’ ss and promote the inclusion of an NMD-inducing exon within intron 49 of the HTT pre-mRNA, leading to down-regulation of mutant HTT mRNA and protein in mouse models of HD. Optimization of the HTT-C2 class of molecules has led to the advancement of these molecules for treatment of HD in clinical trials ^17^.

Although both risdiplam and branaplam enhance the recognition of weak non-canonical GA/GU 5’ ss, they each target a subset of these 5’ ss with preferred upstream and/or downstream sequences. For example, RNAseq analysis of splicing events affected by treatment of cells with either risdiplam or branaplam demonstrated that target 5’ splice sites were highly enriched for the exons with a GA at the -2 and -1 position, with a preference for adenosine at -4 for risdiplam and -3 for branaplan ^17^. Massive parallel splicing assay followed by RNAseq analyses further revealed that the risdiplam class of compounds prefers the ANGA/GUADG (D = U, A, or G, N = any) sequence, while the branaplam class of compounds works on either the above motif or the AGA/GURNG (R = purine) motif ^17,18^. The first motif is present in the 5’ ss of SMN2 intron 7 and the second motif is present in the 5’ ss of a compound-induced Exon (iExon) within HTT intron 49. These added sequence preferences provide impressive target specificity for these compounds. For example, RNA-seq analyses show that branaplam treatment at 24nM (approximately 2 times the IC50 for HTT) for 24 hours only affects 165 out of numerous possible splicing events in human SH-SY5Y cells ^17^.

The molecular mechanism by which these structurally different compounds improve the recognition of the non-canonical GA/GU sequence with additional upstream and downstream sequence specificity is unclear. An NMR structure of short oligos representing the 5’ ss (10 nt) and U1 snRNA (11 nt) indicates that the 5’ ss and U1 snRNA duplex form a bulge where the non-canonical -1A of the weak 5’ ss flips out, likely preventing efficient binding of the 5’ ss to U1 snRNP ^19^. The structure of this short RNA duplex with SMN-C5 (a risdiplam analog) suggests that SMN-C5 brings -1A in and repairs the bulge, strengthening the interaction between the 5’ ss and U1 snRNA ^19^. However, the Kd between the compound and the short RNA duplex (28 µM) is over 1000-fold higher than the EC50 (15 - 30 nM) of the compound in cells ^19,20^, likely due to the lack of any other components of U1 snRNP in the NMR structure. Furthermore, the structure cannot explain the sequence specificity upstream and downstream of the -1A position, as the critical adenosine at the -4 position was not included in the target RNA. In this study, we utilize biochemical approaches with purified native U1 snRNP and reconstituted U1 snRNP, coupled to in vivo assessments of splicing activity, to understand the mechanism of these small molecule splicing modulators, which is critical for developing additional compounds into therapeutics for other splicing-related diseases.

## Results

### Branaplam improves the binding of U1 snRNP to weak 5’ ss in vitro

To evaluate the effect of branaplam on U1 snRNP’s ability to bind weak 5’ ss, we used a fluorescence polarization (FP) experiment with a fluorescently labeled RNA oligo containing sequence around the 5’ ss of exon 7 in the SMN2 pre-mRNA (AGGA/GUAAGUCU, where / indicates the exon and intron boundary, designated as SMN2 5’ ss oligo) or around the 5’ ss of an iExon within intron 49 in the HTT pre-mRNA (CAGA/GUAAGGGU, designated as HTT 5’ ss). We engineered a protein A-tag on U1A in HeLa cells using CRISPR-Cas9 and purified native U1 snRNP using the protein A tag and IgG resin (**Supplemental Fig. 1A**). We titrated increasing concentrations of purified native U1 snRNP and determined the Kd between U1 snRNP and these weak 5’ ss in the presence of DMSO or branaplam. Since U1C is weakly associated with U1 snRNP and may fall off at low U1 snRNP concentrations ^21^, we added a large excess of full-length U1C (800nM) expressed in and purified from E. coli in all binding reactions unless otherwise stated. Using this experiment, we found that branaplam increased the binding affinity between native U1 snRNP and the SMN2 or HTT 5’ ss oligo by about twofold (**Fig. 1A-C**). Although the affinity difference is small, it is highly reproducible.

**Figure 1.**
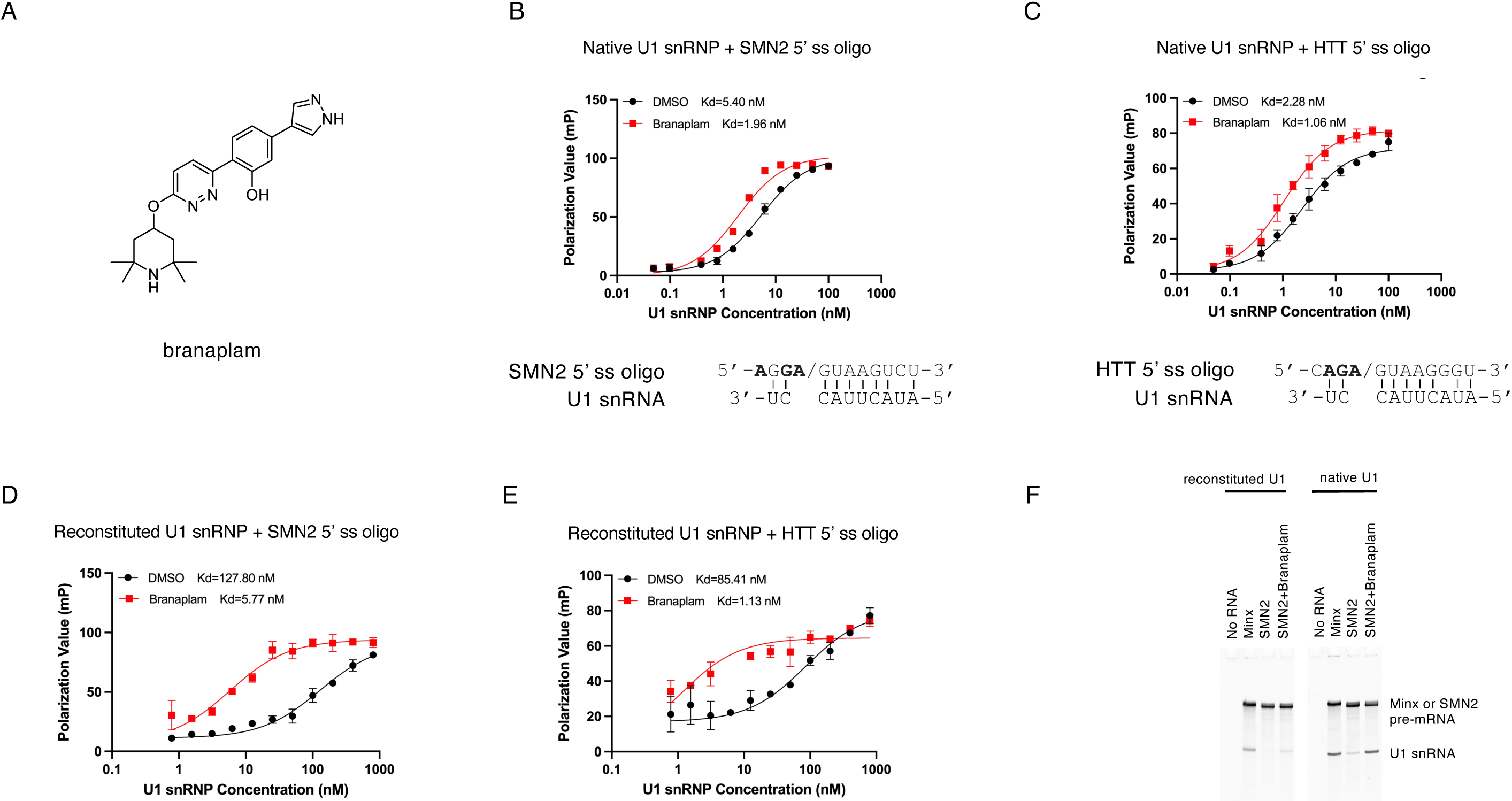
Branaplam improves the binding of native or reconstituted U1 snRNP to both the SMN2 and HTT 5’ ss. A. Structure of branaplam. B. Branaplam improves the binding affinity between native U1 snRNP and the SMN2 5’ ss oligo in fluorescence polarization (FP) experiment. In all FP experiments described in this paper, the 5’ ss oligo was fluorescently labeled, and increasing concentrations of U1 snRNP was added to the binding reaction in the presence of DMSO, or 10 μM branaplam dissolved in the same concentration of DMSO, unless otherwise stated. C. Branaplam improves the binding affinity between native U1 snRNP and the HTT 5’ ss oligo in FP experiment. D. Branaplam improves the binding affinity between reconstituted U1 snRNP and the SMN 5’ ss oligo in FP experiment. E. Branaplam improves the binding affinity between reconstituted U1 snRNP and the HTT 5’ ss oligo in FP experiment. F. MBP bound MS2 tagged Minx (as a positive control with the canonical 5’ ss) or SMN2 pre-mRNA containing the weak 5’ ss of exon 7 were immobilized on amylose resin, incubated with either native or reconstituted U1 snRNP in the absence or presence of branaplam. RNAs from the eluted samples were analyzed on a denaturing gel with SybrGold staining.

To evaluate the effect of branaplam on reconstituted U1 snRNP, protein components were expressed in and purified from E. coli, using the constructs gifted by Dr. Frederick Allain and following essentially their protocol ^22^. Among these, the SmB construct contains residues 1 to 174 (the full length contains 240 residues) and the U1-70K construct contains residues 1-216 (the C-terminal RS domain was truncated). The other subunits are all full-length. We reconstituted the U1 snRNP using these protein components and full-length U1 snRNA and purified the reconstituted U1 snRNP using a superdex 200 column (**Supplemental Fig. 1B**). Similar to the previous result when using purified native U1 snRNP, branaplam dramatically increased the binding affinity between the reconstituted U1 snRNP and the SMN2 or HTT 5’ ss oligo (**Fig. 1D, E**).

As an orthogonal method, we used pulldown assays to evaluate the effect of branaplam on U1 snRNP and 5’ ss binding. In this assay, we immobilized Minx (as a positive control with a canonical 5’ ss) or SMN2 pre-mRNA containing the weak 5’ ss of exon 7 on amylose resin, pulled down either native or reconstituted U1 snRNP, and analyzed the RNAs in the pulldown samples as a surrogate of the amount of U1 snRNP bound to the pre-mRNA. We found that branaplam significantly increases the amount of native or reconstituted U1 snRNPs pulled down by the SMN2 pre-mRNA (**Fig. 1F**). We also found that SMN2 pre-mRNA pulls down more native than reconstituted U1 snRNP in all conditions, suggesting that the native U1 snRNP binds the SMN2 pre-mRNA better than the reconstituted U1 snRNP, consistent with our observations in FP experiments (**Fig. 1B-E**).

### Risdiplam does not improve the binding of U1 snRNP to weak 5’ ss in vitro

We next evaluated the effect of risdiplam on the binding between U1 snRNP and weak 5’ ss containing the GA/GU motif. To our surprise, risdiplam (or its analogs such as SMN-C2) has no effect on the binding between either native or reconstituted U1 snRNP and the SMN2 5’ ss oligo (or the FOXM1 5’ ss oligo with the sequence of **A**U**GA/**GUAAGUUC) (**Fig. 2A-C, Supplemental Fig. 2**). Due to this limitation, all our subsequent in vitro experiments are performed using branaplam and analogs.

**Figure 2.**
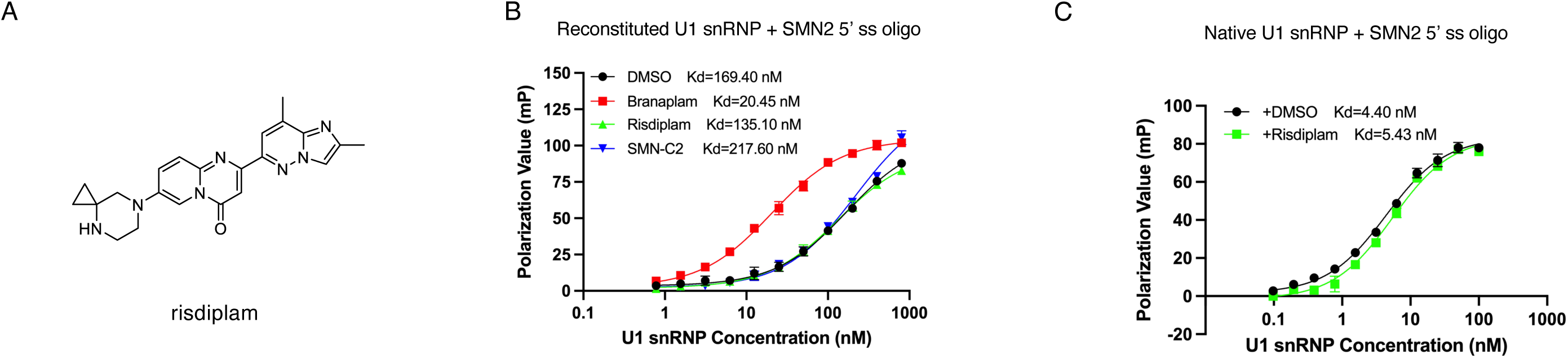
Risdiplam does not improve the binding of U1 snRNP to weak 5’ ss in vitro. A. Structure of risdiplam. B. Branaplam, but not risdiplam or its SMN-C2 analog, improves the binding of reconstituted U1 snRNP to the SMN2 5’ ss oligo in FP experiments. C. Risdiplam has no effect on the binding of native U1 snRNP to the SMN2 5’ ss oligo in FP experiments.

### Branaplam analogs have remarkably similar in vitro and in vivo activity profiles

To assess how our in vitro binding assay correlates with the cellular activity of branaplam analogs, we synthesized a series of branaplam analogs with various substituents on the pyridazine core (**Supplemental Fig. 3**). We evaluated the cellular effect of these analogs on inducing SMN2 exon 7 inclusion using a splicing reporter assay in HEK293 cells and showed that they have various levels of activity with the order: branaplam ∼ SM2 ∼ ZCT-A10 > ZCT-A14 > ZCT-A13 > ZCT-A9 (**Fig. 3A**). We next evaluated the effect of these analogs on the binding affinity between U1 snRNP and the SMN2 or HTT 5’ ss oligo in FP experiments (**Fig. 3B, 3C**). The orders of the in vitro effect of these analogs are remarkably consistent with their in vivo cellular activities in the splicing reporter assay (**Fig. 3D**).

**Figure 3.**
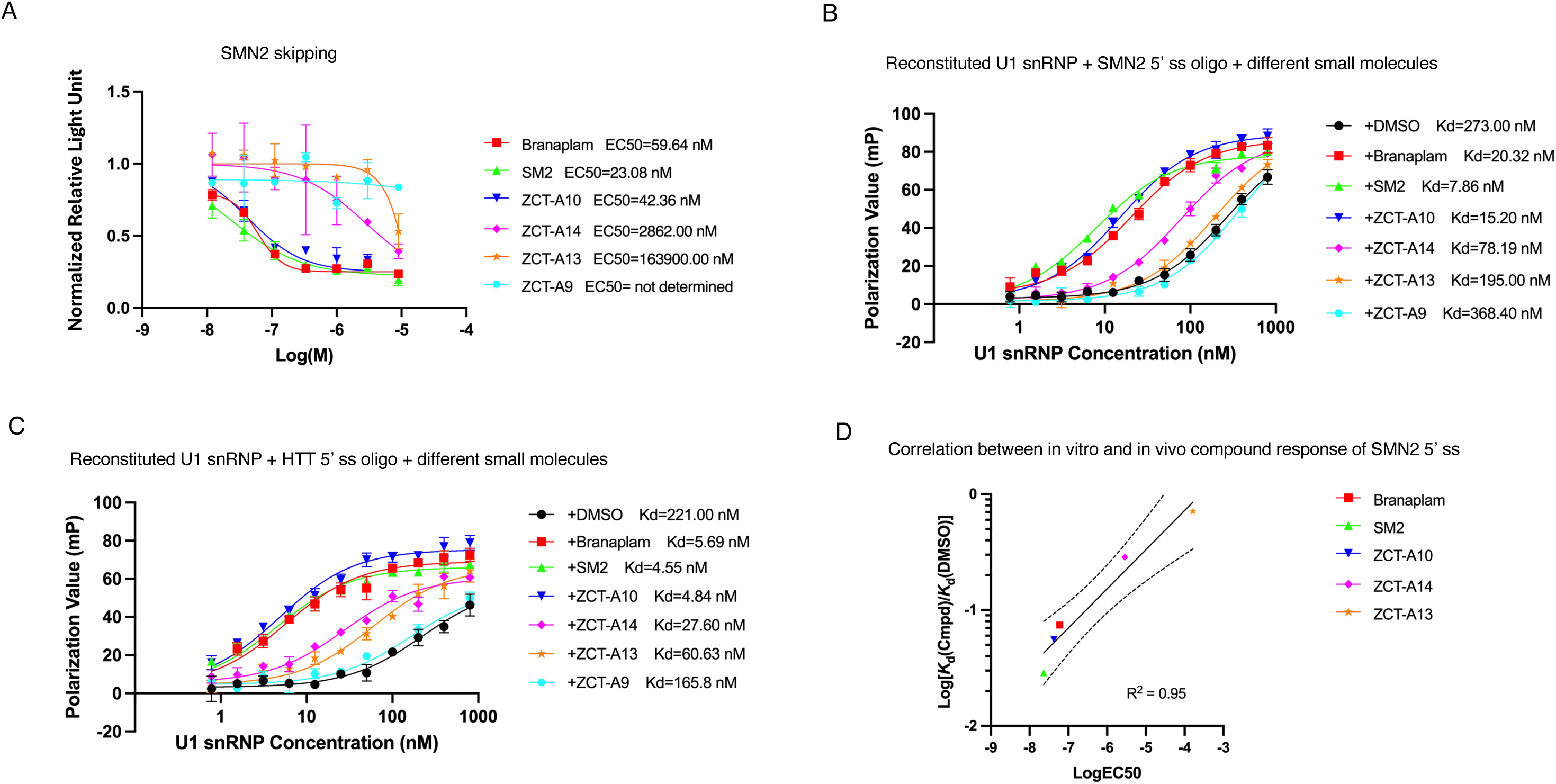
Various branaplam analogs have similar activity profiles in inducing SMN2 exon 7 inclusion in cells and in binding to the SMN2 or HTT 5’ ss oligo in vitro. A. A SMN2 exon7 reporter plasmid was transformed into HEK293T cells, treated with increasing concentrations of branaplam analogs, and the extent of SMN2 exon 7 skipping was evaluated using a luciferase reporter (skipping of exon 7 leads to higher luciferase signal, see Methods for details) and quantified. B. FP experiments were carried out using fluorescently labeled SMN2 5’ ss oligo, increasing concentrations of reconstituted U1 snRNP, in the presence of DMSO or 10 μM of branaplam analogs. C. FP experiments were carried out using fluorescently labeled HTT 5’ ss oligo, increasing concentrations of reconstituted U1 snRNP, in the presence of DMSO or 10 μM of branaplam analogs. D. Scatterplot of log(EC50) values from the reporter assay versus log[Kd(compound)/Kd(DMSO)] from the in vitro binding assay using SMN2 5’ ss, showing a strong correlation between cellular potency and direct binding affinity in vitro. The solid line denotes the best-fit linear regression (R² = 0.95), and the dashed lines represent the 95% confidence interval.

### Mutations of critical nucleotides near the 5’ ss abolish the effect of branaplam

Massive parallel splicing assays (MPSA) revealed that ANGA/GUHDNN and NAGA/GUNNNN are two 5’ splice site motifs responsive to splicing modulators ^18^. The first motif is reflected in the SMN2 5’ ss (**A**G**GA**/GUAAGUCU, referred to as the -4A motif) and the second motif is reflected in the HTT 5’ ss (C**AGA**/GUAAGGGU, referred to as the -3A motif). RNAseq analysis of exons included upon treatment with splicing modulators showed that a distinct specificity for the -3A motif with branaplam and -4A motif with risdiplam ^17^. We mutated nucleotides at several key positions in these motifs and evaluated their effect on U1 snRNP recognition of these 5’ ss in the presence of branaplam. For the HTT oligo, A^-1^U (nucleotide A at -1 position mutated to U), G^-2^U, and A^-3^C mutants all abolish the effect of branaplam (**Fig. 4A-C**). We chose the mutant nucleotide to be different from that in the motif but avoided the canonical 5’ ss nucleotides at these positions (A at -2 and G at -1 positions). For the SMN2 oligo, the A^-1^U and A^-4^C mutants also abolished the effect of branaplam (**Fig. 4D, E**). These results are all consistent with motifs revealed by RNAseq and MPSA analyses, supporting that the in vitro effect of branaplam on reconstituted U1 snRNP and 5’ ss binding recapitulates its effect in cells, thus it is likely that direct compound interaction with the U1-5’ss RNA interface is the primary contributor to the in vivo sequence specificity ^17,18^.

**Figure 4.**
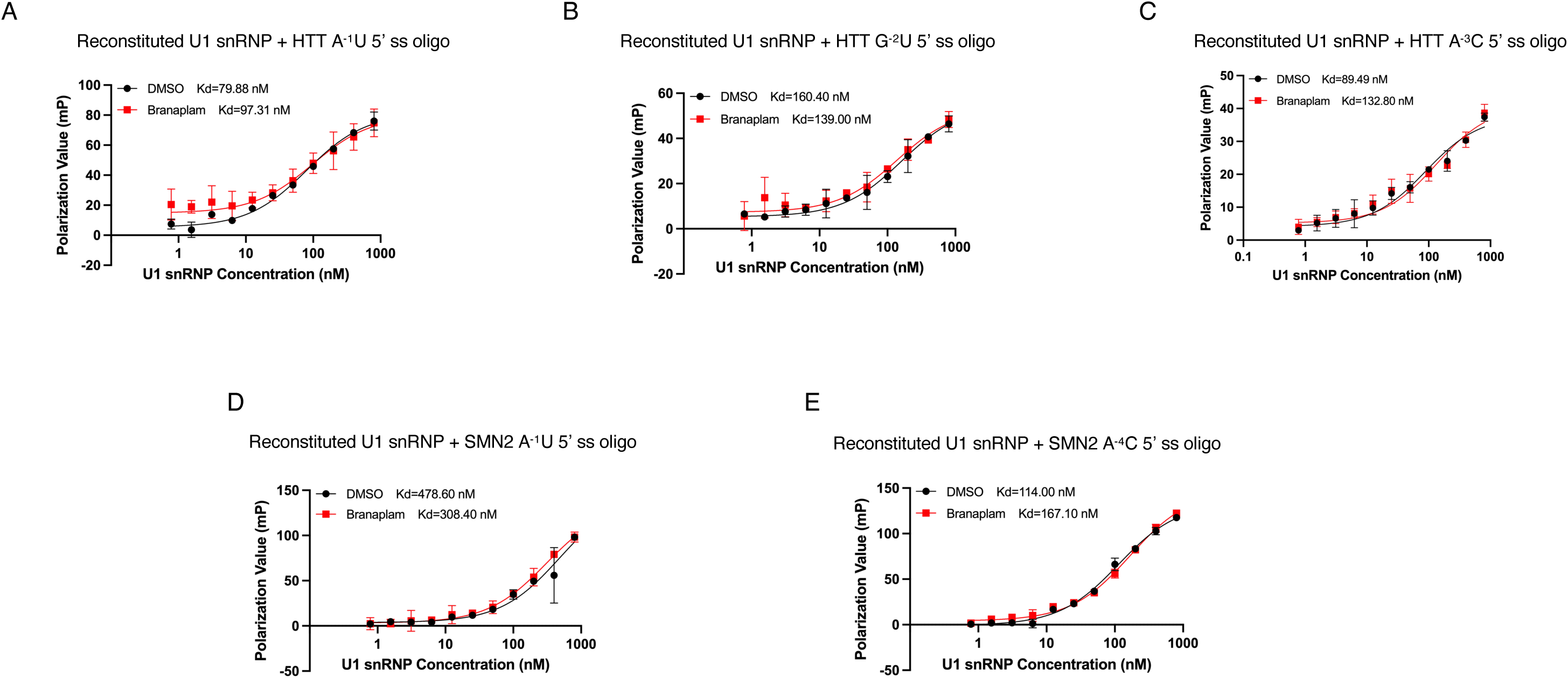
Mutations of critical nucleotides in the SMN2 and HTT 5’ ss oligo abolish the effect of branaplam on 5’ ss binding. FP experiments were performed using fluorescently labeled HTT (A-C) or SMN2 (D-E) oligo carrying the indicated mutant and increasing concentrations of reconstituted U1 snRNP in the presence of DMSO or branaplam.

### U1 snRNA and 5’ ss duplex with the A bulge is not sufficient for the effect of branaplam

We next took advantage of the modular nature of reconstituted U1 snRNP to decipher the components required for branaplam’s effect on U1 snRNP and weak 5’ ss binding. It was speculated that the -1A bulge formed between the 5’ end of U1 snRNA and the weak 5’ ss containing GA/GU is critical for the effect of risdiplam analogs (and potentially branaplam) ^22^. We found that the 5’ end of U1 snRNA (nucleotide 1-10) or the entire U1 snRNA has no substantial binding to the HTT 5’ ss oligo (-4 to +8 around the 5’ ss), even in the presence of U1C (**Supplemental Fig. 4A-C**). We suspected in this in vitro system (with only the 5’ ss oligo, U1 snRNA, and U1C), the bulge A at the -1 position is not actually formed in the U1 snRNA and 5’ ss RNA duplex, potentially due to nucleotides 9 and 10 in U1 snRNA are not forming stable basepairs with nucleotides -2G and -3A of the HTT 5’ ss oligo. Indeed, when we deleted the -1A nucleotide to allow continuous base pairing between the HTT 5’ ss oligo and the U1 snRNA, there is clear binding between the HTT 5’ ss oligo and the U1 snRNA (either the 5’ end or the entire U1 snRNA) (**Supplemental Fig. 4D, E**). As expected, this binding was not improved by branaplam (**Supplemental Fig. 4D, E**), since there is no -1A bulge. To ensure the presence of a - 1A bulge in the RNA duplex, we extended the U1 snRNA to nucleotide 1-16 and generated an artificial HTT 5’ ss oligo that added 4 additional nucleotides upstream of the -4C nucleotide to ensure stable base pairing upstream of the potential -1A bulge. U1 snRNA binds to this artificial HTT 5’ ss oligo, but branaplam has no effect on the binding affinity (**Supplemental Fig. 4F**). These results indicate that the -1A bulge in the U1 snRNA and 5’ ss duplex alone without any U1 snRNP protein component is not sufficient for the effect of branaplam.

### Helix H of U1 snRNP is important for the effect of branaplam

We next evaluated the role of U1 snRNA, especially Helix H (**Fig. 5A**), in the effect of branaplam. To this end, we generated a U1 snRNA tetra mutant (C^119^U/U^120^A/G^121^U/C^122^A) that completely disrupt Helix H. U1 snRNP reconstituted with this snRNA still binds the HTT 5’ ss oligo, but this binding is no longer improved by the presence of branaplam (**Fig. 5B**). We next generated a single G^12^U mutant in Helix H and showed that this mutant also abolishes the effect of branaplam on improving the binding between U1 snRNP and HTT 5’ ss (**Fig. 5C**). Furthermore, when we generated a compensatory mutant to restore Helix H (G^12^U/C^122^A), the effect of branaplam on 5’ ss binding is also restored (**Fig. 5D**), indicating the presence of Helix H in U1 snRNA is important for the effect of branaplam.

**Figure 5.**
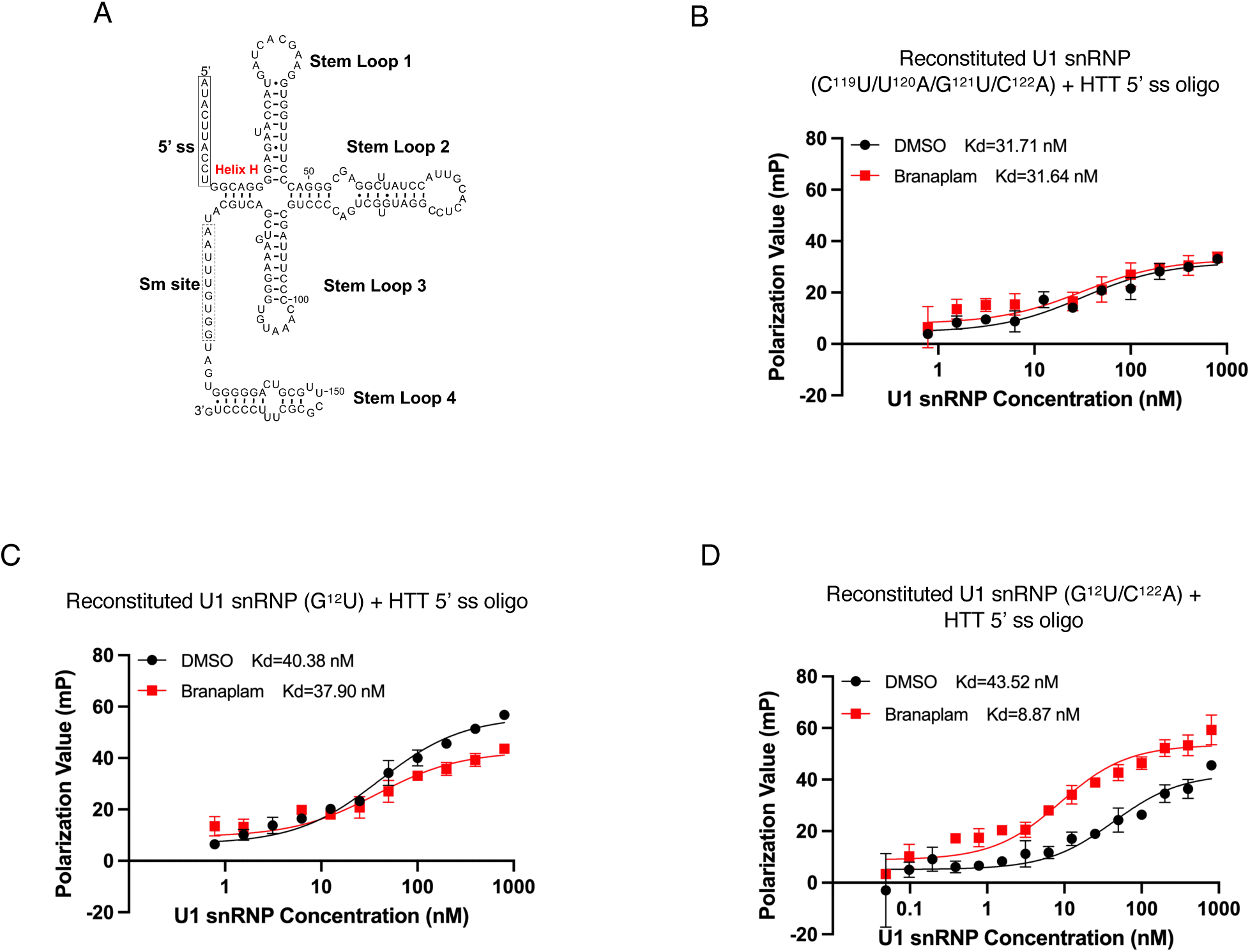
Helix H of U1 snRNA is important for the effect of branaplam on 5’ ss binding in vitro. A. Secondary structure of U1 snRNA indicating the position of Helix H. B. U1 snRNP was reconstituted using a U1 snRNA carrying a tetra-mutant that disrupts Helix H and used in FP experiments. C. U1 snRNP was reconstituted using a U1 snRNA carrying the G12U mutant that disrupts the first basepair of Helix H and used in FP experiments. D. U1 snRNP was reconstituted using a U1 snRNA carrying a double mutant G12U/C122A that restores Helix H and used in FP experiments.

### Helix H is critical for the recognition of 5’ ss in vivo

To understand the importance of Helix H to splicing in vivo, we utilized a U1 variant that has been engineered to recognize the non-canonical GA/GU 5’ splice sites by altering U1 snRNA sequence at position 9 and 10 from CU to UC, complementary to -1G and -2A at the non-canonical 5’ splice site (designated as U1-GA variant). Previous studies demonstrated that this U1 snRNA variant is incorporated into U1 snRNP and targets a set of iExons containing GA/GU 5’ splice site ^17^. When the U1-GA variant is combined with Helix H mutations, we can now measure the in vivo effect of mutations to U1 Helix H on splicing of the exons activated by the U1-GA variant. After expression of either U1-GA variant with wild-type Helix H or U1-GA variant with mutations to key residues in Helix H, we performed RNA sequencing and monitored the inclusion of exons with GA/GU 5’ ss sequences by calculating the deltaPSI (between cells transfected with U1 variants and mock transfected) and determined the effect of mutations to Helix H. Our results showed that mutation of the G12 residue at the base of Helix H (**Fig. 5A**) negatively impacts the ability of the U1-GA variant to function in splicing resulting in loss or diminished splicing of most GA/GU iExons (**Fig. 6, Supplemental Table 1**). Multiple compensatory mutations that restore the base pairing (G^12^A/C^122^U or G^12^C/C^122^G), but alter the identity of the nucleotides, restored the majority of the splicing activity of GA/GU exons in vivo (**Fig. 6, Supplemental Table 1**). The effect of these Helix H mutations on in vivo splicing of GA/GU iExons is consistent with our in vitro observations and suggests a stable Helix H is a key determinant for the recognition of GA/GU 5’ ss by either the U1-GA variant or the compound-mediated mechanism.

**Figure 6.**
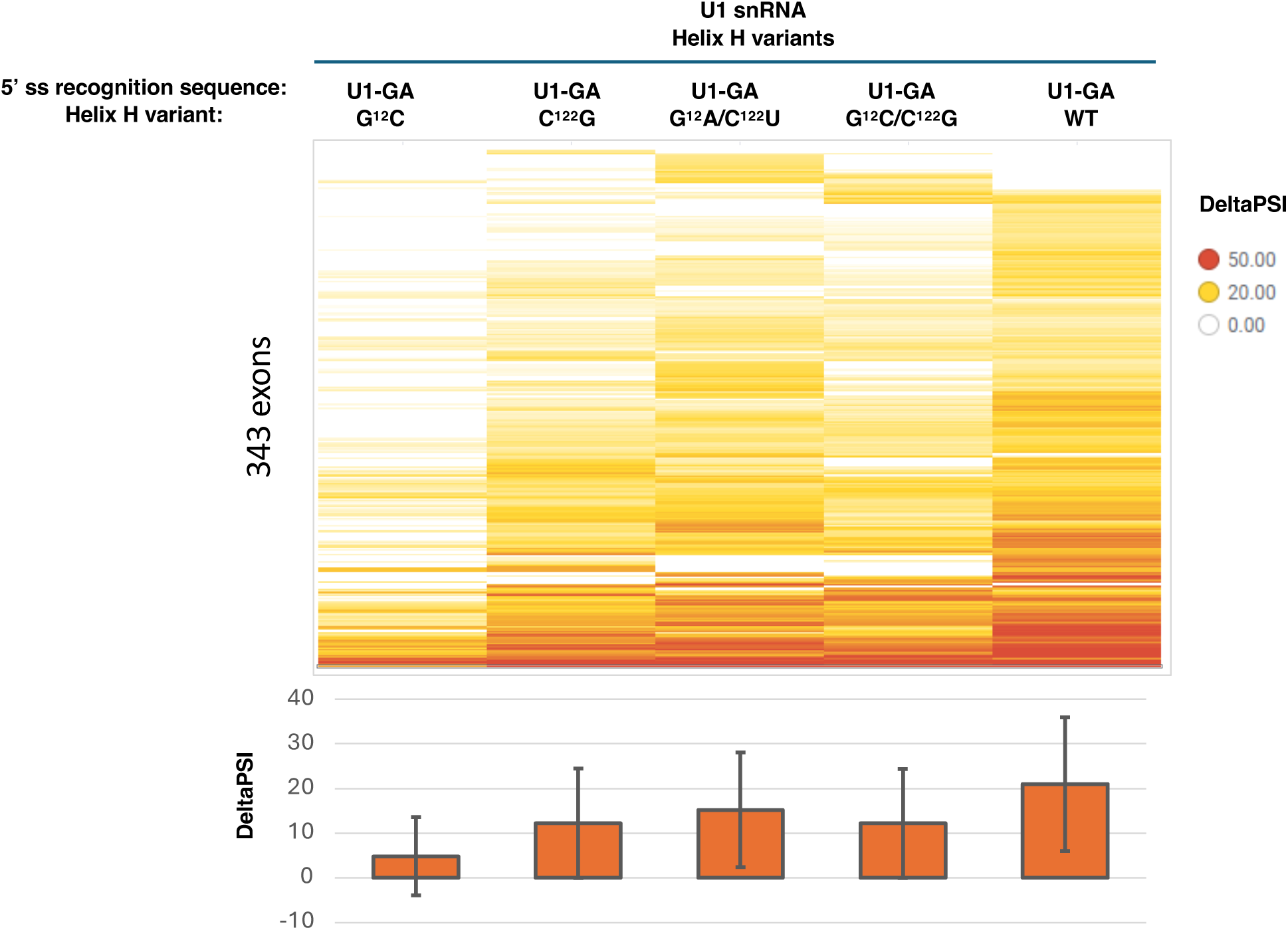
Helix H structure and nucleotide identity are critical to U1 snRNA function in vivo. U1-GA variants each carrying mutations in Helix H were expressed in HEK293 cells and then RNAseq was utilized to determine the level of inclusion of exons containing a GA/GU 5’ splice site. The top figure depicts a heat map of deltaPSI of inclusion events relative to a mock transfection (which contains endogenous U1 without the U1-GA plasmid) for 343 exons which showed inclusion in at least one treatment conditions. The bottom figure depicts a bar plot of the average deltaPSI values for each treatment, with error bars representing the standard deviation.

### U1C but not U1A or U1-70K is important for the effect of branaplam on 5’ ss binding in vitro

We next asked which U1 snRNP protein component may be important for the effect of branaplam. U1 snRNP contains seven Sm proteins that are shared with other snRNPs and three U1 snRNP specific proteins, U1A, U1C, and U1-70K (**Fig. 7A**). We evaluated the role of the three U1 specific proteins in the effect of branaplam. We first reconstituted U1 snRNP without U1A or U1C. We found that branaplam still dramatically improves the binding of U1 snRNP to the HTT 5’ ss oligo in the U1A depleted U1 snRNP but no longer does so for U1 snRNP without U1C (**Fig. 7B, C**). The addition of a truncated U1C (residues 1-61) that contains the ZnF domain without the disordered C-terminal domain (**Fig. 7A**) restores the effect of branaplam (**Fig. 7D**). These data together demonstrate that the ZnF domain of U1C, but not U1A protein, is important for the effect of branaplam on 5’ ss binding in vitro.

**Figure 7.**
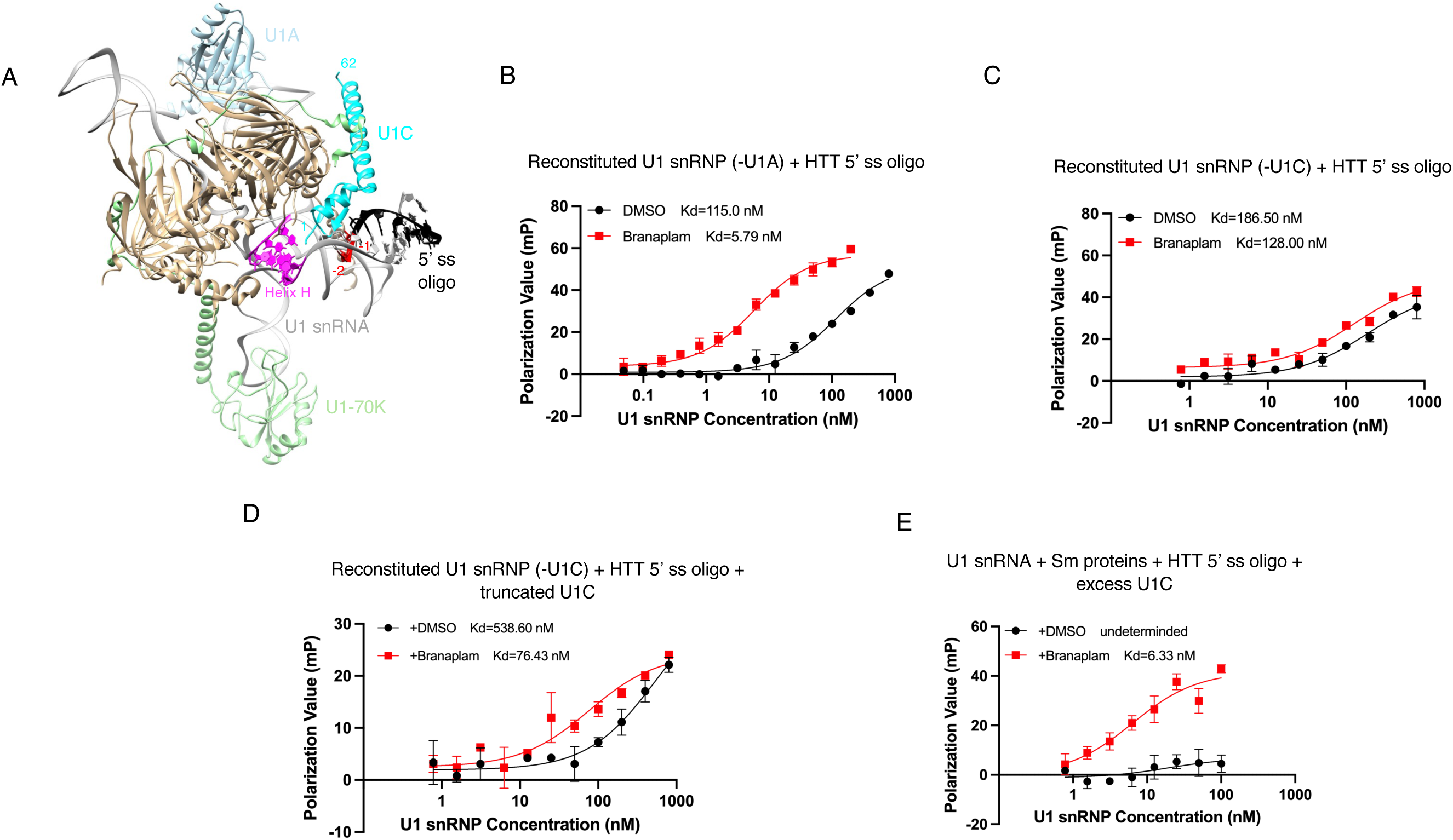
U1C but not U1A or U1-70K is important for the effect of branaplam on 5’ ss binding in vitro. A. A model of reconstituted U1 snRNP based on previous low resolution or partial structures of human U1 snRNPs ^9,10,37^. Black is the 5’ ss oligo used in the crystal structure of a minimal U1 snRNP that fully complement the 5’ end of U1 snRNA ^10^. -1 and -2 positions (relative to the 5’ ss) of the oligo are indicated in red. Purple represent Helix H on the U1 snRNA. Numbers on U1C indicate residue numbers. B. U1 snRNP was reconstituted without U1A and used for FP experiments with fluorescently labeled HTT oligo in the presence of DMSO or branaplam. C. U1 snRNP was reconstituted without U1C and used for FP experiments with fluorescently labeled HTT oligo in the presence of DMSO or branaplam. D. U1 snRNP was reconstituted without U1C and used for FP experiments with fluorescently labeled HTT oligo in the presence of DMSO or branaplam as well as excess truncated U1C (residues 1-61). E. U1 snRNP was reconstituted with U1 snRNA, Sm proteins, and 373 μM of U1C and used for FP experiments with fluorescently labeled HTT oligo in the presence of DMSO or branaplam.

It is more difficult to evaluate the effect of U1-70K depletion because the N-terminus of U1-70K interacts with U1C and the removal of U1-70K will also remove U1C from U1 snRNP (**Fig. 7A**). To overcome this problem, we assembled the U1 snRNA with Sm proteins, then added large excess of U1C protein (373 μM). We showed that branaplam was able to improve the binding of this reconstituted U1 snRNP (without U1A and U1-70K) to the HTT oligo (**Fig. 7E**), suggesting that U1-70K is not required for the effect of branaplam.

### U1C modifies activities of branaplam and risdiplam in vivo

To substantiate the role of U1C in compound-mediated splicing in cells, we knocked down U1C by siRNA in HEK293 cells (**Supplemental Fig. 5**) and treated cells with either risdiplam or branaplam in dose-response for 24 hours. Knockdown of U1C alone had a substantial effect on splicing, leading to skipping of 8364 cassette exons throughout the genome in a sequence-independent manner (**Supplemental Fig. 6A, B, Supplemental Table 2**), consistent with previous published studies of U1C ^23^. Skipped exons are highly enriched for noncanonical 5’ ss sequences, which account for 90% of skipped exons (**Supplemental Fig. 6C**). Notably, among skipped exons with a canonical exonic 5’ss sequence, 74% contain noncanonical intron sequence, consistent with the previous published role for U1C stabilization of the U1 snRNA-pre-mRNA duplex at the 5’ ss ^24–27^ (**Supplemental Fig. 6C, Supplemental Table 2**). Interestingly, U1C knockdown also led to the inclusion of over 750 cassette exons (**Supplemental Fig. 6A**), suggesting that in addition to its role in stabilization of 5’ splice site, U1C also plays a role in suppression of splicing for a subset of exons.

We next examined the effect of U1C knockdown on compound-mediated splicing. Low inclusion exons with GA/GU 5’ splice site that are only detected in response to branaplam or risdiplam display the previously observed sequence preference for A at -3 or -4 position from the splice site and canonical intronic sequence GUAAG at +1 to +6 of the 5’ ss ^17^ (**Supplemental Fig. 7A)**. Comparison of GA/GU splicing events induced by compound show that a reduction in U1C leads to a significant decrease in the potency of both branaplam and risdiplam for a majority of iExon inclusion events (**Fig. 8A, B, Supplemental Fig. 8A-C, Supplemental Fig. 9**). This global effect does not alter the inherent sequence specificity of branaplam and risdiplam at the -4 and -3 position (**Supplemental Fig. 7B**), resulting in the same rank order of potencies for compound effect on GA/GU target exons (**Supplemental Table 2**).

**Figure 8.**
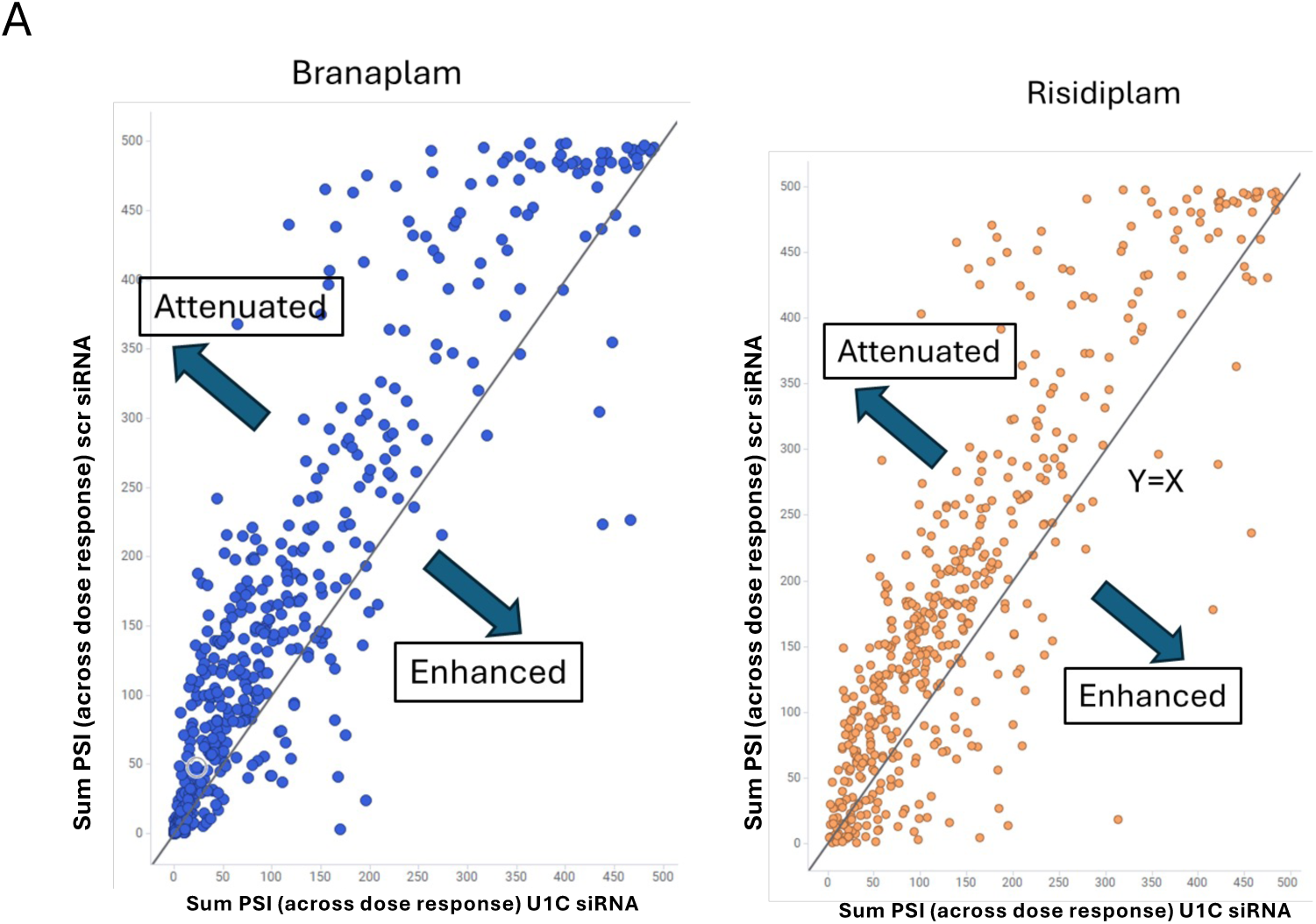
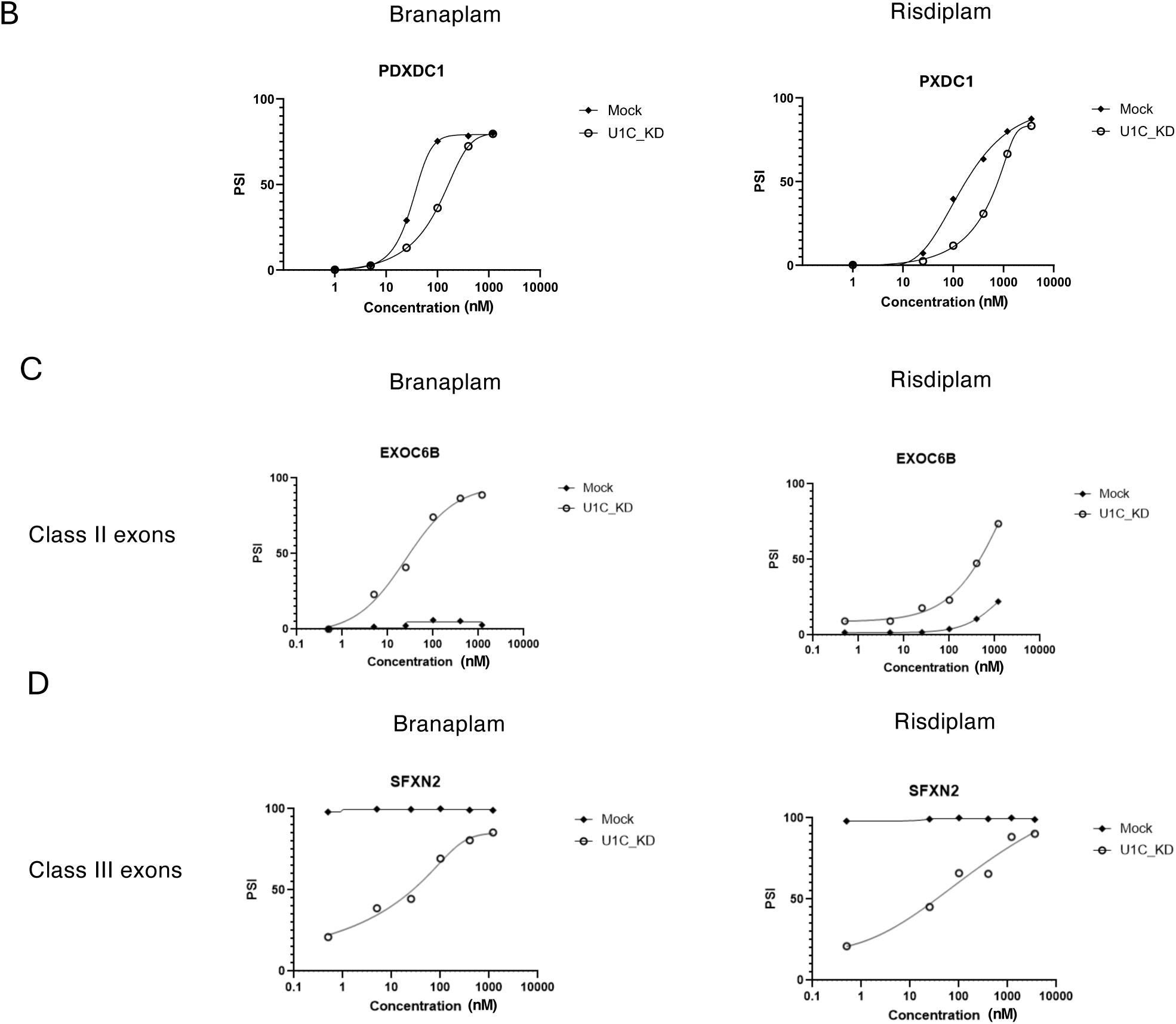
U1C knockdown causes genome-wide exon skipping and alters splicing modulator activity. A. Scatter plot of the sum of PSI for scr (y axis) or U1C (x axis) siRNA treated cells across the dose response of compound (branaplam or risdiplam) for each iExon inclusion event that contains a GA/GU 5’ss. The no effect line (y=x) is plotted. Events where splicing is attenuated shift to the left of the no effect line and those that are enhanced shift to the right of the no effect line. The compound-mediated inclusion of most iExons was attenuated by U1C knockdown. B. Plot of PSI of a representative exon induced by splicing modulator compound treatment. The inclusion of these exons by compound is reduced in U1C knockdown. C. Plot of PSI of a representative class II exon induced by splicing modulator compound treatment, which represents a very low inclusion exon where splicing inclusion by compound is greatly enhance by U1C knockdown. D. Plot of PSI of a representative class III exon induced by splicing modulator compound treatment, where U1C knockdown results in skipping of the exon that is restored when compound is added.

Intriguingly, we also observed a set of low inclusion exons where compound activity is greatly enhanced upon U1C knockdown (**Fig. 8C, Supplemental Fig. 8D**). In most cases, both branaplam and risdiplam demonstrate a similar increase in potency, while for other targets such as PPARG (**Supplemental Fig. 8D**), only branaplam activity is enhanced. Additionally, of the 8364 exons skipped after U1C KD, 158 contain GA/GU 5’ ss, 78 of which are at least partially rescued at the highest dose of either risdiplam or branaplam (**Fig. 8D, Supplemental Fig. 8E**). Combined with the observation that U1C knockdown alone is sufficient to cause inclusion of many cassette exons (**Supplemental Fig. 6A, B**), this suggests that U1C may suppress splicing of a subset of exons in a context dependent manner and once U1C levels are reduced, the activity of compounds that promote inclusion of these exons can be demonstrated.

## Discussion

To understand the mechanism of how small molecule splicing modulators improve the recognition of weak 5’ ss with an AGA/GU or ANGA/GU sequence by U1 snRNP, we first evaluated the effect of branaplam on the binding affinity between purified native U1 snRNP and either a SMN2 or HTT 5’ ss oligo. We showed that branaplam consistently increased binding affinity by 2-3 folds (**Fig. 1B, C, F**). We next showed that branaplam also dramatically increases the binding affinity between either the SMN2 or HTT 5’ ss oligo and U1 snRNP reconstituted from protein components purified from *E. coli* and in vitro transcribed U1 snRNA (**Fig. 1D, E, F**). We synthesized a number of branaplam analogs with various degrees of activity in inhibiting SMN2 exon 7 skipping in cells (**Fig. 3A**). The activity profiles of these compounds for improving the binding of U1 snRNP to either the SMN2 or HTT 5’ ss are strikingly similar to their cellular activities in preventing SMN2 exon 7 skipping (**Fig. 3B, C**). Ishigami and colleagues speculated that branaplam binds to the SMN2 and HTT motifs in two very different modes ^18^. However, the similar activity profile of the branaplam analogs for both the SMN2 and HTT 5’ ss suggests that branaplam binds to both motifs using a similar mode.

We further showed that when the A^-1^, G^-2^, and A^-3^ nucleotide in the HTT oligo, or A^-1^ and A^-4^ nucleotide in the SMN2 oligo are mutated, branaplam no longer increases the binding affinity between reconstituted U1 snRNP and these 5’ ss oligos (**Fig. 3**). These observations are consistent with the branaplam responsive sequence motifs observed in cells using massive parallel splicing assays and RNAseq experiments ^17,18^, indicating that the reconstituted U1 snRNP contains all key elements required for branaplam’s effect on U1-5’ ss binding. The failure of a mutated HTT or SMN2 5’ splice with U^-1^ bulge to respond to branaplam demonstrates that the nucleotide identity at the -1 bulge is a key determinant for branaplam recognition (**Fig. 4A, D**).

To further understand the determinants for branaplam function, we took advantage of the modular nature of the reconstituted U1 snRNP. We first showed that a U1 snRNA and HTT 5’ss duplex alone is not sufficient for the branaplam effect (**Supplemental Fig. 2**). We next showed that Helix H of U1 snRNA is important for branaplam binding (**Fig. 5**). Disrupting the first set of base pairs (G12-C122) in Helix H abolished the effect of branaplam. Restoring this to a (U12-A122) basepair restores the response to branaplam, suggesting that the basepair nature of Helix H is important for the effect of branaplam. White and colleagues demonstrated that a U1 snRNP containing a minimal U1 snRNA construct (60 nt in length, with Helix H maintained but most of the stem loops 1-3 beyond Helix H replaced by a kissing loop) modified from the one used for crystallographic studies in Kiyoshi Nagai’s lab ^10^ also responds to branaplam ^21^, supporting the importance of Helix H in compound response. We further evaluated the effect of Helix H mutants on GA/GU 5’ ss recognition by U1 snRNP in vivo. Utilizing a U1-GA variant, we demonstrated that compensatory changes that maintain base pairing in Helix H (G^12^A-C^122^U or G^12^C-C^122^G) are functional, establishing a critical role for base pairing at the base of Helix H in vivo. Interestingly, there is a significantly larger effect on loss of splicing with the G^12^C mutant than with the C^122^G mutant, suggesting that the identity of a guanosine at position 12 is also important for the proper function of U1 snRNA in vivo. Together, the in vitro and in vivo data demonstrate that U1 snRNP recognition of the GA/GU 5’ ss and compound engagement both require a specific structural configuration of Helix H.

We next demonstrated that the ZnF domain of U1C but not U1A or U1-70K is also a determinant for the branaplam function (**Fig. 7**), consistent with previous observation by White and colleagues that U1C but not U1A is needed for the effect of branaplam in minimal U1 snRNP ^21^. U1C has been proposed to stabilize the interaction of weak 5’ splice sites to U1 snRNP through direct interaction with the RNA duplex formed upon U1 snRNP binding ^10^. We considered two possible roles for U1C in splicing modulator function. The first is that U1C functions to promote a conformation among the ensemble of possible structures of the U1 snRNA and 5’ ss duplex that is conducive to compound binding, similar to what has been proposed in other systems ^28,29^. The second possibility, as suggested by White et al., is that compound binding depends on direct interaction with U1C which is required for modulation of U1-GA/GU 5’ss recognition ^21^.

To gain transcriptome-wide insight into these two potential models for U1C’s role in compound activity, we tested the effect of U1C KD in vivo on splicing modulator potency by RNA-seq. Our data demonstrated that U1C knockdown broadly reduces the effect of branaplam and risdiplam (**Fig. 8B, Supplemental Fig. 8A-C, Supplemental Fig. 9**). While the respective sequence specificities of branaplam and risdiplam do not change upon U1C knockdown, the majority of compound-induced GA/GU iExon inclusion events display 2-3-fold decrease in potency, consistent with our in vitro observation on the importance of U1C in the effect of branaplam. This reduction is not as dramatic as our in vitro observations, likely because some U1C remains in U1 snRNP (possibly due to a slow turnover rate) 48 hours after U1C KD (**Supplemental Fig. 5**).

The RNA-seq results further revealed three additional classes of exons that provide unique insights into the role of U1C in the effect of splicing modulators. Class I is defined by a set of U1C-suppressed cassette exons wherein U1C KD increases exon inclusion in the absence of compound (comparing U1C KD and scramble control in DMSO, **Supplemental Fig. 6A, B**). These cassette exons do not display any bias for exonic 5’ ss sequences and are never responsive to compound. In this class of events, the presence of U1C could be enhancing the stability of structures that are not conducive to 5’ ss recognition and progression toward splicing. Class II is identified by cassette exons which have GA/GU 5’ splice sites and are either fully skipped or included at very low levels and do not respond to compound treatment when U1C expression is normal. Only upon reduction of U1C expression do these exons become responsive to compound treatment leading to their inclusion (**Fig. 8C**). Class III consists of a subset of the >8000 cassette exons that are skipped when U1C expression is decreased. When exons from this class contain a GA/GU 5’ ss, both branaplam and risdiplam can restore the inclusion of the exon to the level observed in samples with endogenous U1C expression (**Fig. 8D**). Thus, splicing modulator activity occurs at a reduced level of U1C for this class of exons, suggesting splicing modulators can function independent of U1C to promote exon inclusion in certain context.

Taken together, the identification of these three classes of exons suggests that U1C can suppress inclusion of a subset of exons in a context-dependent manner. One possible scenario is that the interaction between U1 snRNA and these 5’ ss is too tight in the presence of U1C, which will hinder subsequent steps in the splicing cycle (such as the removal of U1 snRNP and transition of 5’ ss base pairing to U6 snRNA). The reduction of U1C can potentially release its suppression of exon inclusion in response to compound treatment. However, these exons are mostly noncanonical, and the binding affinity is likely suboptimal, making this scenario unlikely. Although the exact mechanism of U1C-mediated suppression of these subsets of exons remains unclear, the fact that two distinct chemical classes of splicing modulators are active on several classes of exons only upon U1C reduction makes it unlikely that direct contact between U1C and compounds is a driving force of compound activity. Instead, our data are consistent with a model whereby the main role for U1C is to stabilize a specific structural conformation at the interface between U1 snRNA and a GA/GU 5’ss, such as those found in SMN2 or HTT which can be targeted by the compound. The noncanonical nature of targeted exons, where intronic sequence is the canonical GUAAG and exonic sequence is a noncanonical GA/GU sequence, strongly suggests that U1C promotes stabilization within the noncanonical NAGA and ANGA exonic portion of the U1snRNA-5’ss complex. In this model, Helix H, positioned in close proximity to the exonic portion of the 5’ss, likely provides additional stability, as changes that alter the stability abrogate the ability of the compound to bind a U1-5’ss in vitro. Similar changes that alter the stability of Helix H are detrimental to exon inclusion in vivo, demonstrating the importance of structural conformation within this region for 5’ ss recognition. In the context of the class II and III exons, we hypothesize that U1C can decrease the prevalence of the conformation recognized by branaplam and risdiplam which therefore allows for compound activity to be enhanced only upon knockdown of U1C. Understanding the context-dependent effects of 5’ ss sequence and splicing factors on RNA structures available for small molecule binding to enhance 5’ ss usage is critical for the further development of splicing-modulating therapeutics to treat human diseases and the development of RNA-targeting small molecules in general.

Finally, the lack of activity of risdiplam on either the native or reconstituted U1 snRNP’s ability to bind SMN2 5’ ss in vitro, even in the presence of U1C (**Fig. 2A-C**), suggests that additional cellular factors may be needed for the risdiplam effect. One such candidate is the Luc7L family of alternative splicing factors (Luc7L, L2, and L3). These proteins are homologs of the yeast Luc7 protein which binds to the U1 snRNA and 5’ ss RNA duplex on the opposite side of U1C and stabilizes the duplex together with U1C ^30,31^. The presence of exons whose response to branaplam is strongly enhanced, but whose response to risdiplam remains unchanged upon U1C KD (**Supplemental Fig. 8D**), is consistent with the requirement of other cellular factors for the effect of risdiplam, which may render these exons less sensitive to U1C KD. It is possible that additional cellular factors such as Luc7L proteins could contribute to the binding of risdiplam in a similar manner as U1C, by altering the ensemble of structures formed to present a more favorable binding site for the compound.

## Methods

### Engineering HTP tag on endogenous U1A in HeLa cells

Parental Hela-S3 cell lines used in this study were from Cell Technologies Shared Resource of the University of Colorado Anschutz Medical Campus. HeLa-S3 cells were routinely cultured adherently in MEM medium (Corning) supplemented with 10% FBS and 1x penicillin-streptomycin (Invitrogen).

To generate C-terminal fusions of a HTP (HA-His10-TEV-protA) tag in-frame with U1A gene, two guide RNAs (gRNAs) target close to the stop codon of U1A were selected using the gRNA design tool CRISPOR (https://crispor.gi.ucsc.edu/). Each gRNA was inserted into pX330 as described by Ran et al ^32^. Linear doner DNA fragments were generated by PCR using primers U1ADF/U1ADR to incorporate 100bp up and downstream of the stop codon of U1A as the homology arms to an HTP-T2A-neo cassette (the template plasmid pAC5-HTP-T2A-neo was a gift from David Bentley ^33^). Two phosphorothioate linkages were included at the 5’ end of each primer. Hela-S3 cell culturing adherently in 60mm dish were co-transfected with 2 μg each of pX330 gRNA plasmid and linear donor DNA using Lipofectamine LTX (Thermo Fisher Scientific). G418 (800ug/mL) was added 72 h after transfection and the medium was changed every 2–3 days until the G418-sensitive cells were eliminated, and then G418 concentration was dropped to 400ug/ml. After 2-3 weeks, colonies were transferred to 24-well plates, and genomic DNA was isolated for PCR genotyping. For the U1A-HTP colonies, U1AendF /U1A3UTRR (363 bp product) were used to screen for the presence of the wild-type allele, and U1AendF/HTPR (757 bp product) were used to screen for incorporation of the HTP-T2A-neo cassette. The PCR products were confirmed by Sanger sequencing. Western blots were performed using anti-HA antibody (clone 3F10, Roche) to further confirm the fusion of the tag to U1A.

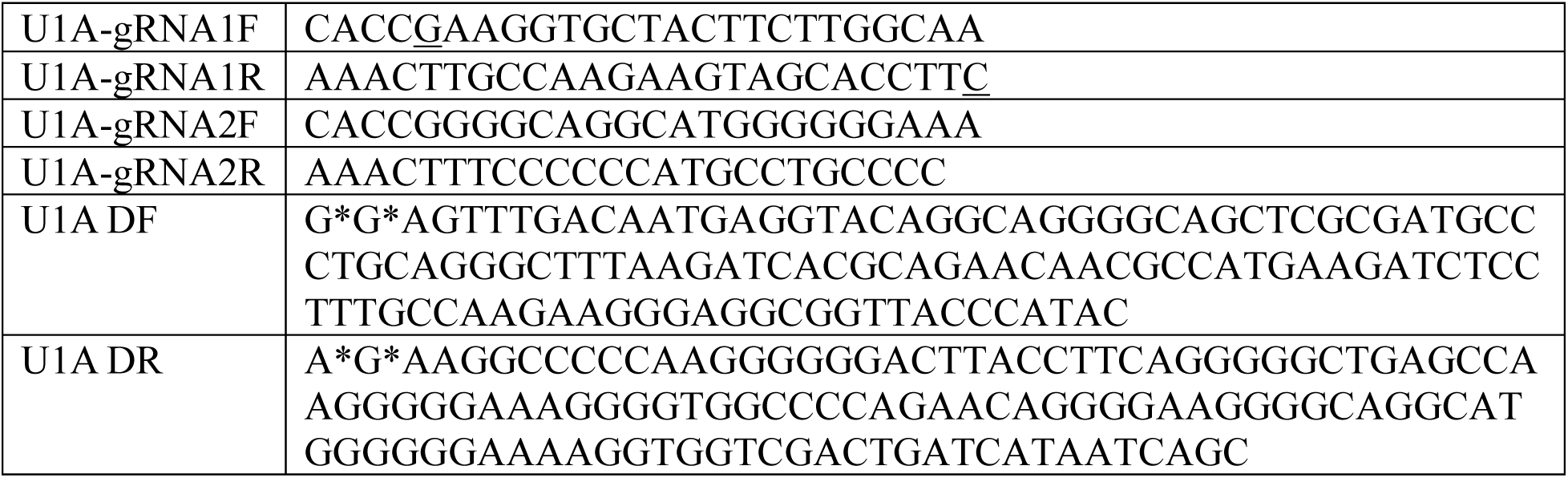
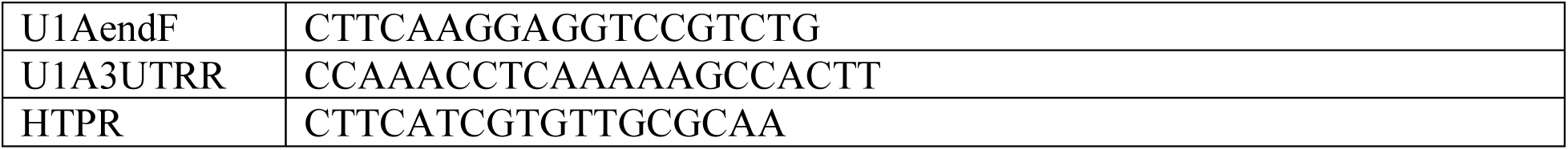

### Native U1 snRNP purification

U1A-HTP cells were adapted to grow in suspension in Joklik’s MEM (Sigma) supplemented with 5% (v/v) newborn calf serum and 1x Pluronic™ F-68 (Gibco). Cells were cultured in 2-liler flasks in an orbital shaker (∼110rpm) at 37°C, and maintained at the density 0.4-1x 109 cells/L tightly. 8-10 liters of cells at density of ∼1x 109 cells/L were harvest and washed with PBS. The cell pellet was snap-frozen for native U1 snRNP purification.

The cell pellet was thaw on ice and gently resuspend in 5 pellet volumes of cold Buffer A (10mM Hepes pH 7.9, 10mM KCl, 1.5mM MgCl2, 0.5mM DTT) and leave on ice for 10 min. The cell suspension was centrifuged for 20 min at 25,000 g at 4°C. The supernatant was removed. The nuclei pellet was Resuspend in cold Buffer IP 250 (20mM Hepes pH 7.9, 250mM NaCl, 1.5mM MgCl2, 0.05% NP-40, 0.5mM DTT) with protease inhibitors (Sigma), and sonicated 2-3 times for 10 second at 40% amplitude. The lysate was centrifuge for 30min at 25,000g and the supernatant was incubated with 0.5-1ml of preequilibrated IgG Sepharose 6 affinity resins (Cytiva) for 2-3hr. The resins were then washed with buffer IP 250, IP 200 (same with IP 250 with 200mM NaCl), IP 150 (same with IP 250 with 150mM NaCl) without NP-40. The U1 snRNP was released by TEV cleavage O/N at 4C. The purified U1 snRNP was further concentrated using an Amicon centrifugal concentrator, and analyzed by SDS-PAGE and RNA gel for the presence all the U1 snRNP proteins and U1 snRNA.

### Pull down experiment

The SMN2 pre-mRNA fragment used in pull down experiment contains 100nt up and downstream of the 5’ss of SMN2 intron 7 followed by three copies of MS2 tag. It was cloned downstream of a T7 promoter in the Sp72 vector. The SMN2 pre-mRNA was in vitro transcribed and purified by 7M Urea denaturing gel. 0.5 μM of SMN2-MS2 (pre-incubated with 50-fold excess of MBP-MS2 protein) was incubated with 0.3 μM of reconstituted or native U1 snRNP in buffer (20 mM Hepes, pH 7.5, 150mM KCl, 3mM MaCl_2_, 1 mM TCEP), in presence of 50 μM of Branaplam or 0.5% DMSO. The complex was then incubated with amylose resin and washed with the same buffer. The resins were treated in 5 μl buffer containing 1mg/ml proteinase K at 37 °C for 30 min to release the RNAs. The resulting samples were run on 6% denaturing polyacrylamide gel and stained with Sybr Gold to visualize the RNAs.

### Expression and purification of individual U1 snRNP protein components

The plasmids used for individual protein components of human U1 snRNP expression are gifts kindly provided by Dr. Frédéric Allain ^22^. These include Sm B (1-174)-D3, Sm D1-D2 and Sm E-F-G, U1A, U1C, and U1-70K (1-216). *Escherichia coli* BL21 (DE3) (Novagen) strains transformed with the plasmids above-mentioned using standard procedures were grown to an OD_600_ of ∼0.6 at 37 °C and then induced with 0.5 mM isopropyl β-D-thiogalactoside (IPTG) for further 20 hours at 16 °C.

Cells expressing these proteins were harvested by centrifugation and lysed by sonication, then the individual components were purified to homogeneity by consecutive/sequential immobilized Ni-NTA metal ion affinity chromatography (Qiagen), cation-exchange chromatography (SP Sepharose Fast Flow; GE Healthcare) and gel filtration (Superdex 75 columns, GE Healthcare) chromatography, following the protocol described by Allain and colleagues ^22^. Purified protein was concentrated using a Millipore concentrator and then stored at −80°C.

### *In vitro* transcription and purification of U1 snRNA

The DNA template for full-length U1 snRNA was cloned into the pSP72 plasmid following a T7 promoter. The recombinant vector was digested by PstI restriction endonuclease for linearization at 37°C overnight and subsequently purified using phenol : chloroform extraction followed by ethanol precipitation. Large-scale *in vitro* transcription was performed in 40 μM MgCl2 at 37°C for 3 h. The transcription mixture was applied to denaturing urea polyacrylamide gel electrophoresis (PAGE) and total RNA was extracted, purified using phenol : chloroform extraction followed by ethanol precipitation, dissolved in DEPC-dd H2O and finally stored at - 80°C. Site-specific U1 snRNA mutants were generated by PCR-based method and verified by DNA sequencing. Purification procedure of mutant U1 snRNAs were identical to that of the wild-type U1 snRNA.

### In vitro reconstitution of U1 snRNP

Prior to complex reconstitution, U1 snRNA was first heated at 80°C for 3 min, and then cooling immediately on ice for 10 min. The annealed U1 snRNA was then incubated with Sm B-D3, Sm D1-D2, Sm E-F-G in a molar ratio of 1:1.5:1.5:1.5. After incubation at 30°C for 30 min and at 37°C for 15 min, the mixture was cooled on ice for 5 min, U1–70K (1-216) and U1A in equal molar amounts to U1 snRNA were added successively, with an interval of 20 min. After incubated on ice overnight, U1 snRNP ΔU1C was purified by anion exchange chromatography on a 1 ml RESOURCE^TM^ Q (Cytiva) previously equilibrated with buffer A and eluted with a linear gradient of 0.25–2.0 M KCl though buffer U (10 mM Hepes pH 7.5, 2M KCl, 5 mM DTT). The obtained U1 snRNP ΔU1-C was then dialyzed overnight against buffer V (10 mM sodium phosphate pH6.8, 100 mM NaCl, 5 mM DTT) and calculated the amount. Finally, 2-fold molar amounts of U1-C was then added. After incubated on ice for 4 h, the mixture separated by size exclusion chromatography using a Superdex 200 increase in buffer C. Peak fractions containing the desired U1 snRNP complex were collected and concentrated.

### Fluorescence Polarization Assays

All of 5’-FAM labeled RNA probes used in this study were synthesized and purified by IDT (USA). Fluorescence polarization (FP) assays were performed in binding buffer (20 mM HEPES pH 7.5, 1 mM MgCl2, 200 mM KCl, 0.01% TritonX-100, 0.1%PEG3350, 2%DMSO) at room temperature using a Envision HTS Plate Reader system (Revvity). Each reaction system contains 3 nM of FAM-labeled RNA oligo, reconstituted U1 snRNP (or native U1 snRNP, or full-length U1 snRNA, or 5’end of U1 snRNA) in 2-fold increasing concentrations, 800 nM (for reconstituted U1 snRNP) or 400 nM (for native U1 snRNP) U1C, different small molecules in DMSO (10 μM unless otherwise stated). In some experiments, 10 μM LU7L2 was incubated in dark for 30 min at room temperature. The reported values are from the mean values (±standard deviation, SD) of two independent experiments. The equilibrium dissociation constants (Kd) of binding reaction were calculated according to the following a single-site binding model formula using GraphPad Prism:

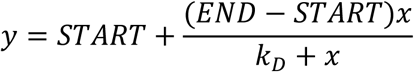

where x and y represent the concentration of U1 snRNP (or U1 snRNA) and the polarization value measured respectively, START represents the polarization value of protein-free nucleic acid probe, END-START represents the maximum polarization value when U1 snRNP (or U1 snRNA) binds to the fluorescently labeled 5’ ss oligo.

### Compound synthesis

See supplemental information.

### SMN2 Splicing Minigene Reporter Assay

The SMN2 splicing reporter minigene plasmid (pCI-SMN2-skipping-GLuc, Addgene #218670) encodes truncated exons 6 and 8, mutated exon 7 (with a single nucleotide insertion), and truncated introns 6 and 7 of the human *SMN2* gene fused to Gaussia luciferase (GLuc), as previously described ^34,35^. Exon 7 inclusion places the GLuc gene out of frame, thereby reducing luciferase activity. HEK293T cells were cultured at 37 °C with 5% CO_2_ in complete growth medium containing 10% FBS (Cytiva, #SH3039603HI) in DMEM (Thermo, #11995073) supplemented with Antibiotic-Antimycotic (Thermo, #15240062). Transfection was performed in 10 cm dishes using Lipofectamine 2000 (Thermo, #11668019) according to the manufacturer’s protocol. 6–16 hours post-transfection, cells were trypsinized (Thermo, #12605010) and seeded into 384-well plates (Greiner, #781080) that had been preloaded with compounds via acoustic dispensing. Compounds were dispensed in duplicate across a 1:3 serial dilution series ranging from 0.012 to 9 µM. Following a 36 h incubation at 37 °C, luciferase activity was assessed by adding 12 µL of a custom luciferase reagent to each well. The luciferase reagent (12 mL total volume) consisted of 60 µL 50× coelenterazine substrate (Nanolight, #320), 3 mL Nanolight GLuc buffer, 30 µL 10% Triton X-100 (Thermo, #A16046.AP), and 8.9 mL deionized water. Plates were shaken at 600 rpm for 6 minutes, and luminescence was measured using a Cytation 5 multimode plate reader (BioTek). Half-maximal effective concentration (EC_50_) values were calculated using GraphPad Prism (v10.4) with four-parameter nonlinear regression curve fitting.

### HelixH U1-GA variants and RNA sequencing

Synthesis of the U1 variants constructs, growth of HEK293 cells, transfection and RNA isolation were described earlier^17^.

### siRNA knockdown of U1C and RNA sequencing

HEK293T cells were grown in DMEM media with 10% FBS. 500,000 cells in 1.75mL of media were plated into a 6 well plate for approximately 6 hours. Either siRNA negative control (ThermoFisher Silencer^TM^ Negative control #2, AM4613) or U1C siRNA (ThermoFisher Silencer^TM^ Select Pre-designed siRNA, S13225) were transfected for a final concentration of 25nM using Invitrogen Lipofectamine RNAiMax (13778-150). In brief, 10 uL 5uM siRNA was added to 115 uL of OptiMEM while 5 uL Lipofectamine added to 120 uL OptiMEM for 5 minutes before mixing the solutions and incubating for 20 minutes. 250uL of Lipofectamine:siRNA solution was gentle added to the 1.75mL cells. After 24 hours post-transfection, cells were treated with 10uL compound resuspended in DMSO. Final concentrations of branaplam were 1.2mM, 400nM, 100nM, 25nM, and 5nM. Final concentrations of risdiplam were 3.6mM, 1.2mM, 400nM, 200nM, and 50nM. Each compound treated experimental condition was treated and collected in duplicate while four DMSO only treated samples per siRNA transfection were collected. RNA was isolated using KingFisher^TM^ Apex instrument and the MagMAX™ *mir*Vana™ Total RNA Isolation Kit (A27828) by Thermo Fisher according to manufacturer’s protocol. Isolated RNA was submitted to Admera Health for library preparation (polyA selected, stranded) and mRNA-sequencing and >20M 150bp pair-end reads were returned.

### RT-qPCR validation of U1C knock down

BioRad iTaq universal SYBR green reaction mix (Bio-Rad 172-5150) was used according to manufacturer’s protocol with modifications. For a total reaction volume of 10 uL, 50 ng of total RNA was amplified with U1C forward and reverse primers at a final concentration of 300nM. Reactions were initial incubated at 50°C for 10 minutes followed by heat inactivation of RT at 95°C for 1 minute. Two-step PCR amplification was performed at 95°C for 10 seconds and 60°C for 30 seconds for 40 cycles followed by measurement of a melt curve. PPIA primers were used to measure an endogenous control and fold change was calculated using ΔΔCt method.

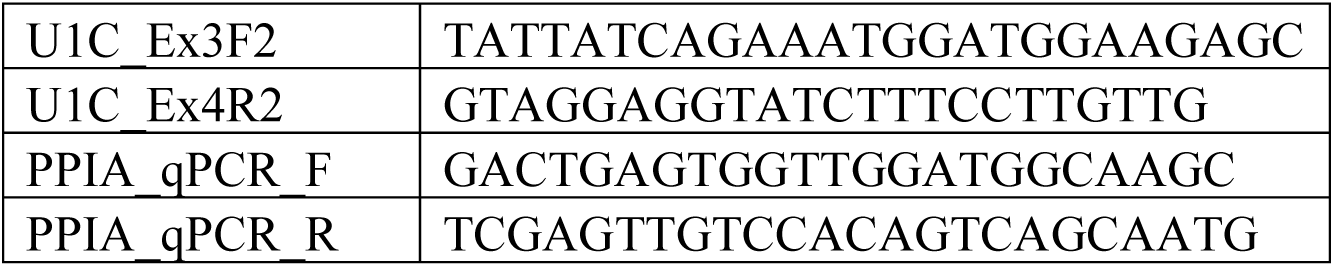

### Evaluation of U1C level in whole cell lysate and U1 snRNP after U1C knock down

Transfection of siU1C for knockdown in HEK293 was performed similarly as for RNA sequencing experiments above. RNA was extracted with a Qiagen RNeasy Plus Mini kit (74134) after lysate homogenization with a Qiashredder (79656). RT-qPCR validation of U1C knockdown is also similar as above, except for use of the New England Biolabs Luna One-Step RT-qPCR kit (E3005S). For protein studies, 48 h cell pellets were lysed with 50 mM Tris-HCl, pH 7.5, 150 mM NaCl, 1% Igepal, 0.25% deoxycholate, 0.1% SDS, and 1X protease inhibitor cocktail (P3100-010) and sonicated with a Diogenode Bioruptor Pico for 2 s x 3 cycles with 1 m rest on ice between cycles. Lysates were clarified by centrifugation at 18,000 g for 20 m at 4°C. For immunoprecipitation, lysate was incubated with U1A antibody (Santa Cruz Biotechnology sc-376027) for 2.5 h at 4°C, followed by incubation with Protein G beads (10004D) 1 h at 4°C. IP samples were washed 3 times with IP Wash Buffer (50 mM Tris-HCl, pH 7.5, 150 mM NaCl, 0.1% Igepal), and eluted with 2X Laemmli sample buffer followed by boiling for 5 m. Whole cell lysates and IP samples were immunoblotted with anti-U1C (Santa Cruz Biotechnology sc-101549), anti-U1A (Santa Cruz Biotechnology sc-376027), and anti-SmB (Y12, gift from Dr. Joan Steitz’s lab^36^).

### RNAseq processing and splicing analysis

RNAseq and splicing analysis were described earlier using DEDseq (https://github.com/liwc01/DEDSeq^17^) with minor modifications. RNA sequencing reads were mapped to human genome (hg19) using STAR (version 2.7.10b); only uniquely mapped reads (with MAPQ > 10) with <5nt/100nt mismatches and properly paired reads were used. For splicing analysis, all junction reads (read with a gap in alignment indicating splicing) were used, including the ones mapped to unannotated splice sites. Reads were counted for different exons (for cassette exon [CE]) or exonic regions (for A5′ss or A3′ss). For each splicing event, a percent-spliced-in (PSI) value was calculated using the percent of average read number supporting the inclusion among all reads supporting either the inclusion or the exclusion. A minimum of 10 for the denominator of PSI calculation was required. Otherwise, a ‘NA’ value would be generated. PSI values for biological replicates were averaged, and the PSI difference between the two treatment groups was calculated. For a statistical test, a 2 × 2 read counts table was made for each event, with rows for reads supporting inclusion or exclusion, and columns for the two comparing sample groups. Fisher’s exact test was used for the statistical tests. For biological replicates, each pair of the testing and control sample was analyzed by Fisher’s exact test and the geometric mean of all *P*-values was reported as a final *P*-value. PSI change (deltaPSI) of >10% (or < −10%) and *P* < 0.001 was used to select splicing events being regulated by the treatment. PSI calculation, Fisher’s exact test, were performed using R (4.0.5). 5’ ss composition analysis was performed as describe in Kenny et al. 2025 ^24^.

## Supplemental Figure Legends

**Supplemental Fig. 1.**
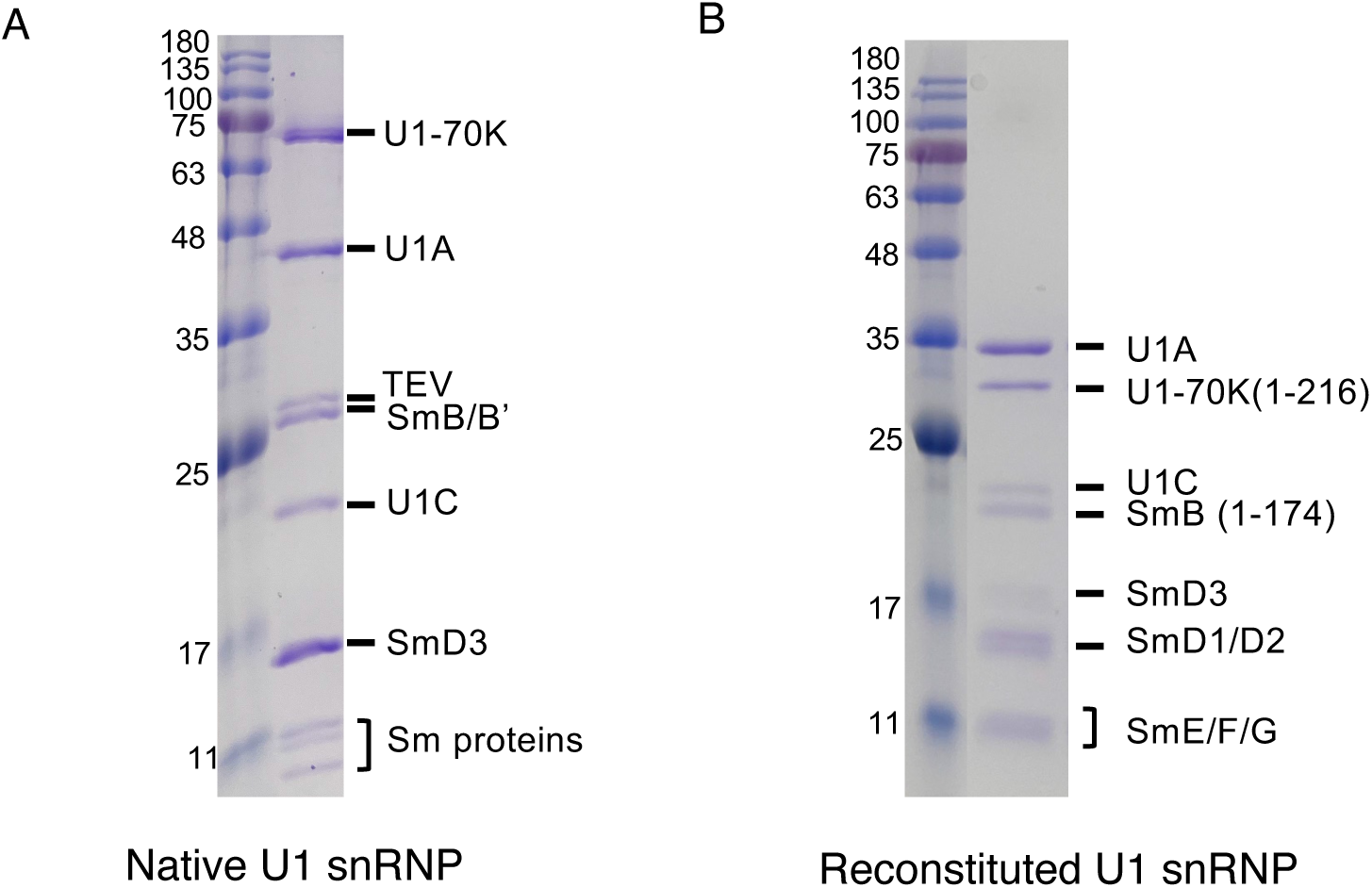
Expression and purification of the native and reconstituted U1 snRNP. A. Coomassie stained SDS PAGE gel of native U1 snRNP purified from HeLa cells using the protein A tag on U1A. B. Coomassie stained SDS PAGE gel of U1 snRNP reconstituted from protein components purified from E. coli.

**Supplemental Fig. 2.**
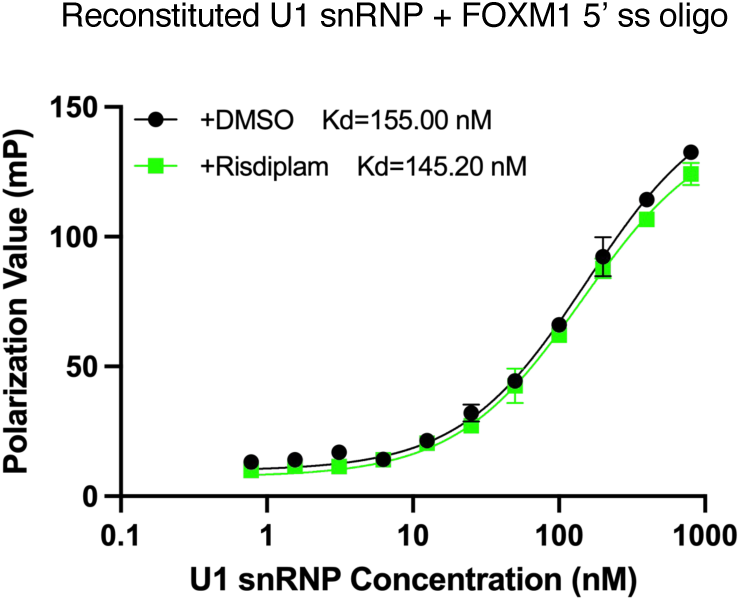
**FP experiments using the FOXM1 5’ ss oligo (AUGA/GUAAGUUC) indicate that risdiplam does not improve the binding of reconstituted U1 snRNP to this oligo that shares the HTT motif (in bold).**

**Supplemental Fig. 3.**
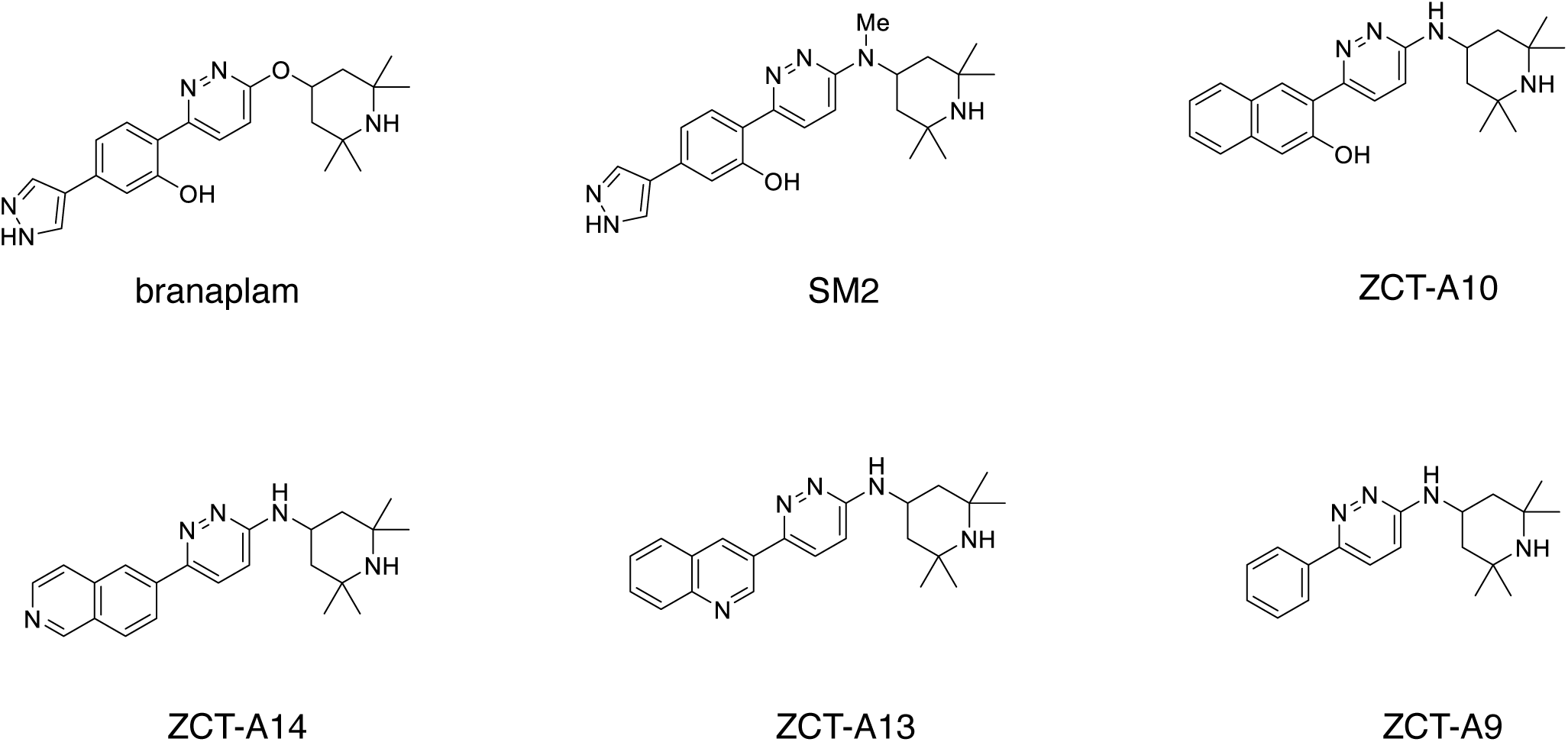
**Structures of branaplam analogs.**

**Supplemental Fig. 4.**
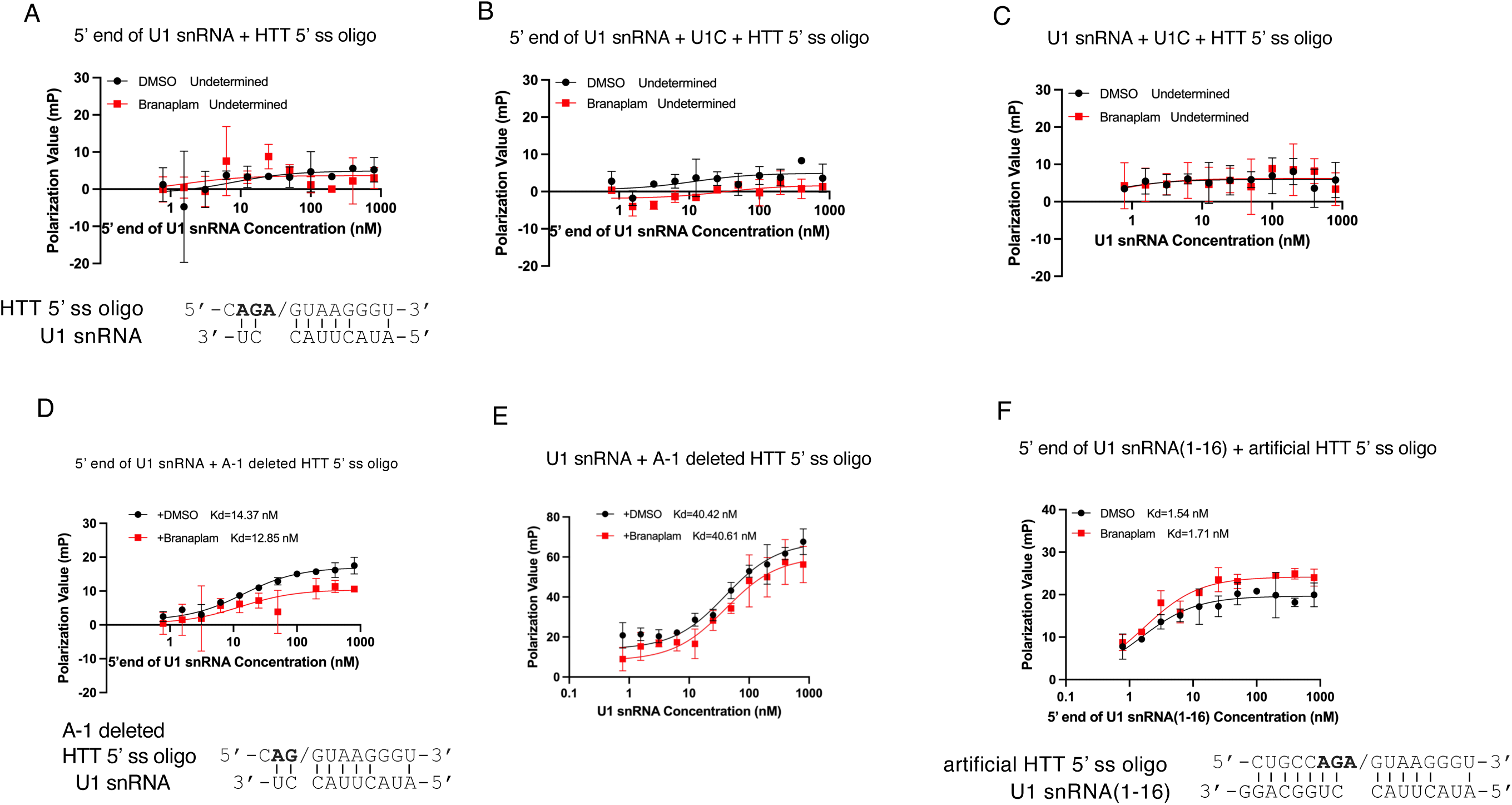
U1 snRNA and 5’ ss duplex with the A bulge is not sufficient for the effect of branaplam. A. FP experiment using fluorescently labeled HTT 5’ ss oligo and increasing concentration of an oligo representing the 5’ end of U1 snRNA (nucleotide 1-10) indicates there is no obvious binding between the two oligos. B. The same FP experiment carried out as in A with the addition of 800 nM of U1C indicates there is no obvious binding between the two oligos even in the presence of U1C. C. FP experiment carried out with the full length instead of the 5’ end of U1 snRNA and the SMN2 5’ ss oligo indicates there is no obvious binding between U1 snRNA and the HTT 5’ ss oligo even in the presence of U1C. D. FP experiment carried out with the -1A nucleotide in the HTT 5’ ss oligo deleted indicates that the removal of the -1A bulge generate appreciable binding between the mutant HTT 5’ ss oligo and the 5’ end of U1 snRNA, but this binding is not improved by the presence of branaplam. E. The same FP experiment was carried out as in D with the full length instead of the 5’ end of U1 snRNA, demonstrating even more substantial binding with the mutant HTT 5’ ss oligo (with the -1A bulge removed), but this binding is not improved by the presence of branaplam. F. An artificial HTT oligo was designed to basepair with nucleotides 1-16 of U1 snRNA and ensure the presence of a -1A bulge. FP experiments using this oligo with a fluorescent label and nucleotide 1-16 of U1 snRNA indicate that these two oligos bind to each other but this binding is not improved by the presence of branaplam.

**Supplemental Fig. 5.**
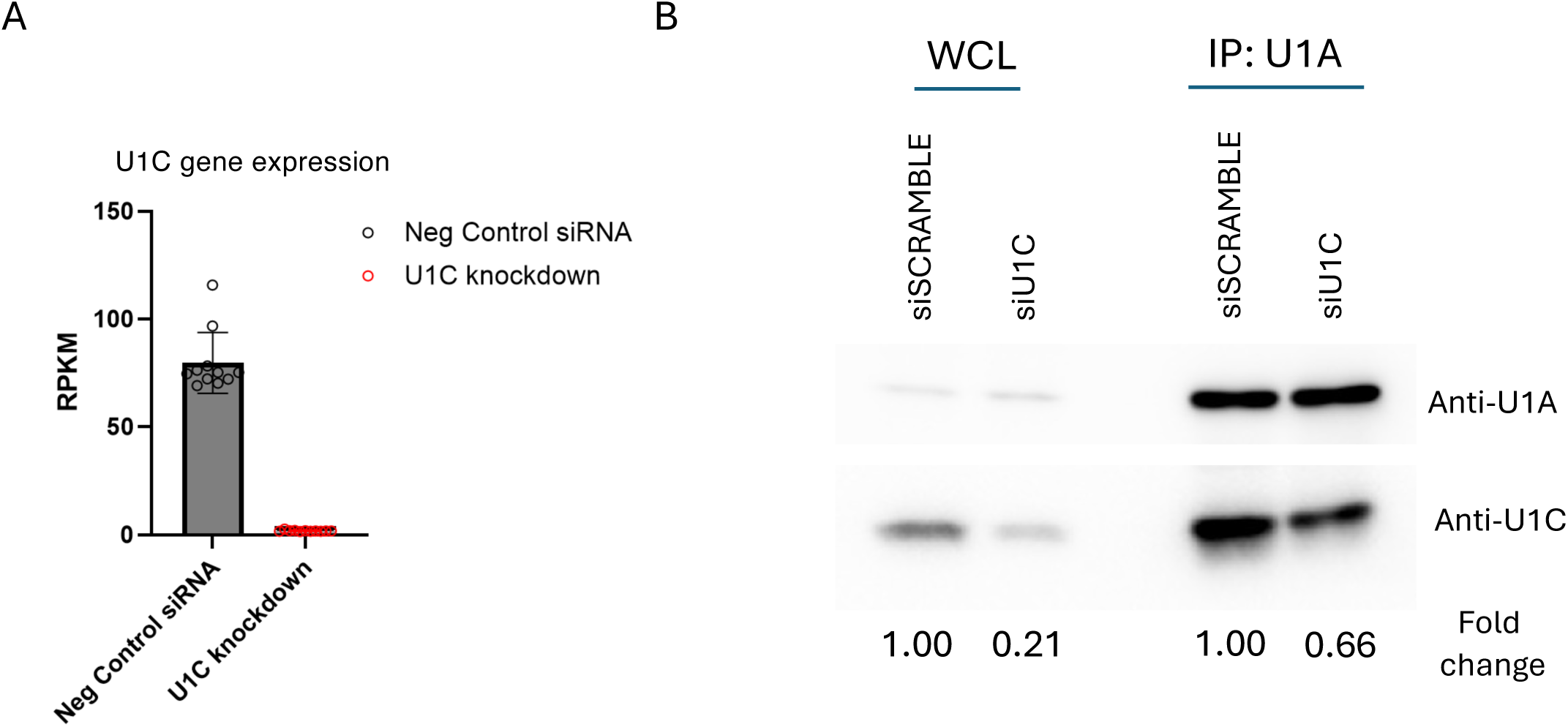
U1C protein level is dramatically reduced in whole cell lysate but retained at a substantial level in U1 snRNP after U1C KD. A. U1C mRNA level determined by qRT-PCR after U1C KD. B. U1C and U1A protein levels in whole cell lysate (WCL) and U1 snRNP (immunoprecipitated by anti-U1A antibody) determined by Western blot.

**Supplemental Fig. 6.**
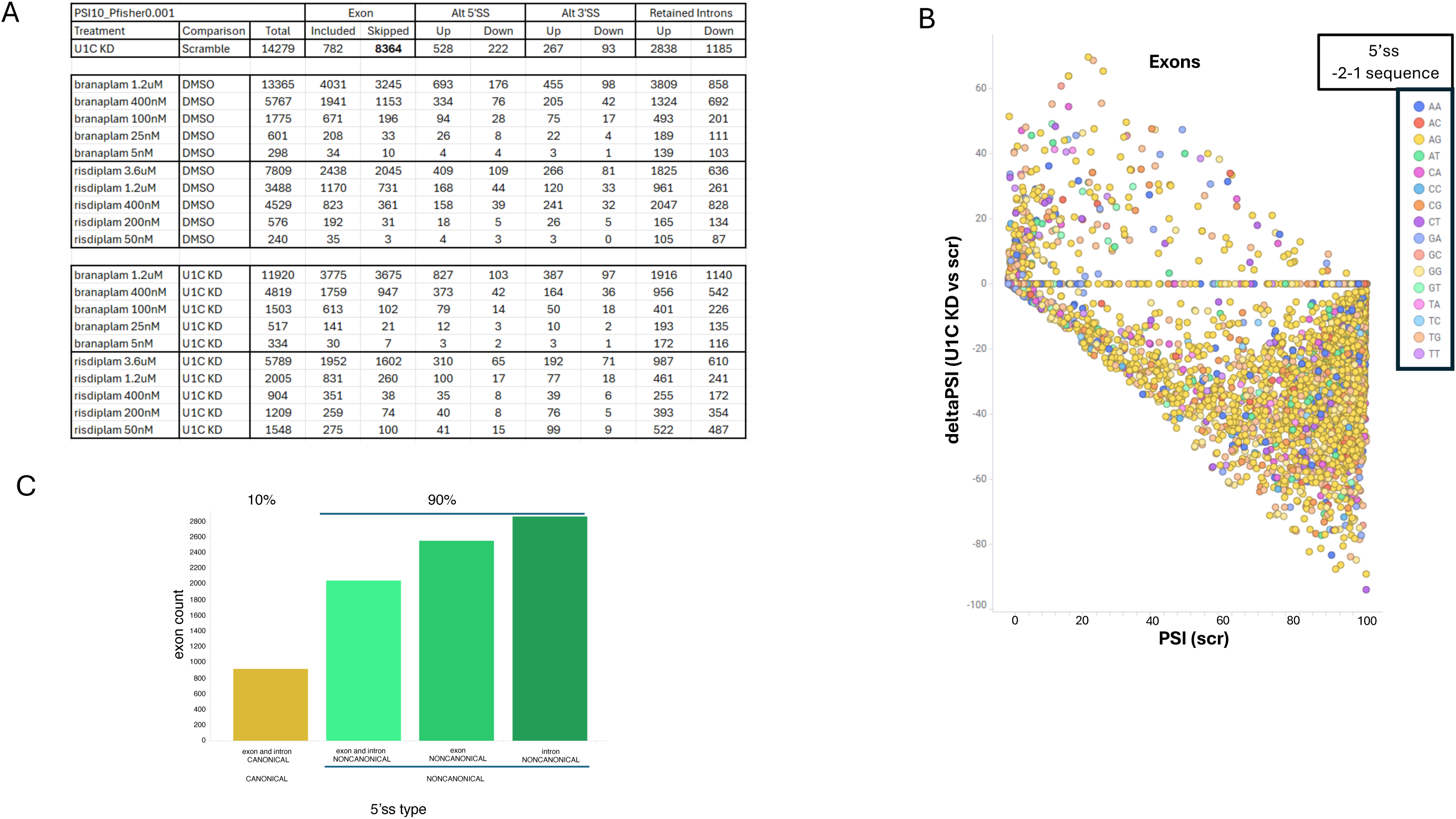
Splicing statistics for U1C Knockdown. A. A summary table of all splicing changes for U1C Knockdown, including statistics of identified exons (included and skipped), Alternative 5’ splice site (Alt 5’SS), Alternative 3’ splice site (Alt 3’SS) and Retained Introns. Row 1, U1C siRNA Knockdown compared to negative control siRNA; Row 4-8 and 9-13, dose response branaplam or risdiplam, normalized to negative control siRNA; Row 15-19 and 20-24, dose response branaplam or risidiplam normalized to U1C siRNA knockdown. B. Scatter plot of cassette exon deltaPSI for all U1C siRNA Knockdown deltaPSI (y-axis) versus PSI of negative control siRNA (baseline PSI; x-axis). Color represents the different -2 and -1 nucleotides at the 5’ss of the exons. C. Bar graph representation of analysis of 5’ ss sequence composition of skipped exons, indicating enrichment for noncanonical 5’ ss. Canonical and noncanonical nature of the exonic and intron portions of the 5’ ss were calculated as described by Kenny et.al, 2025 ^24^ for each skipped exon (n=8364) upon U1C KD. If both exon or intron score is canonical, exon is considered CANONICAL; if either or both scores are noncanonical, exon is considered NONCANONICAL.

**Supplemental Fig. 7.**
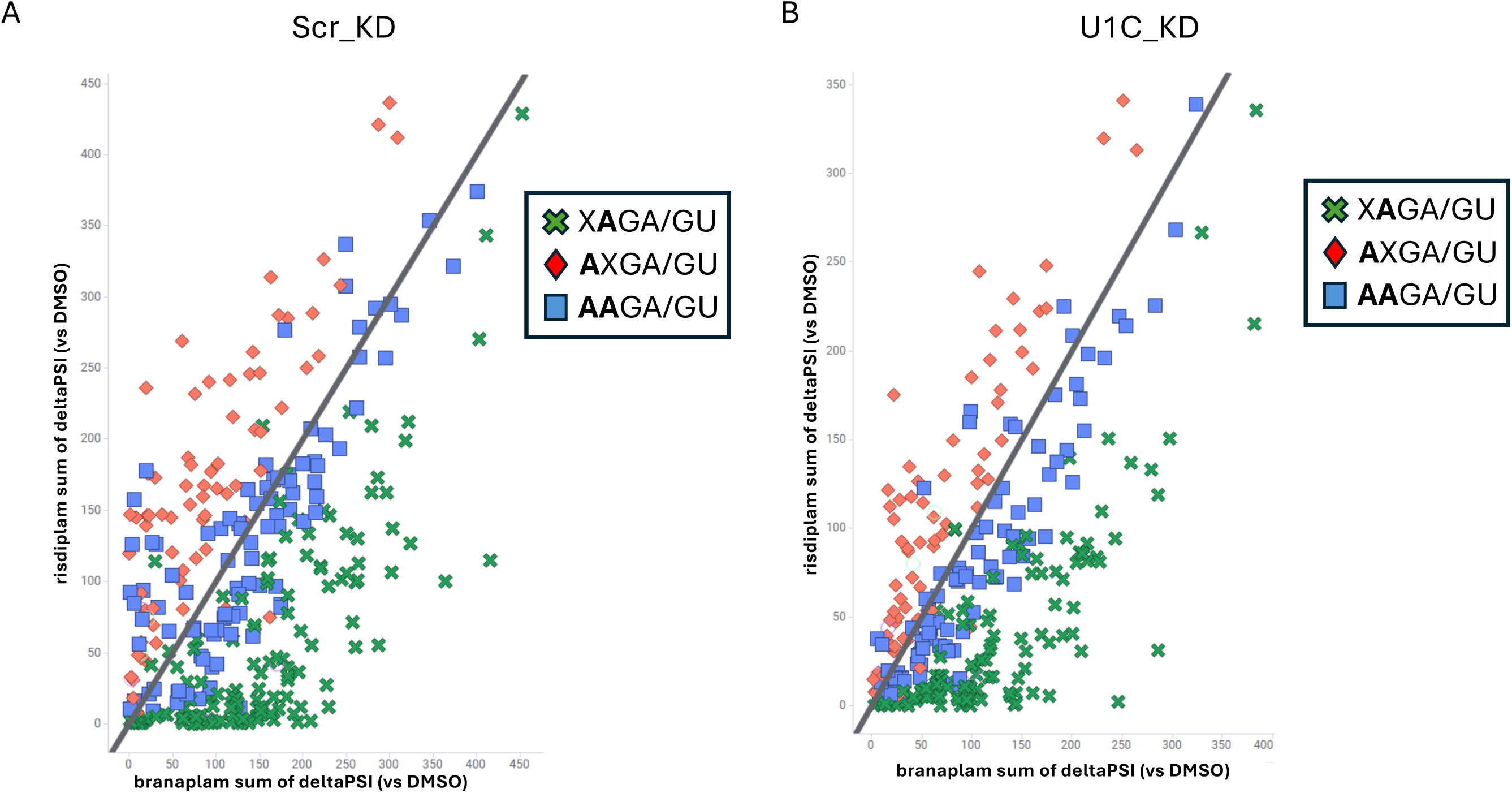
U1C knock down does not affect the sequence preference for either branaplam or risdiplam. A. Scatter plot of Risdiplam sum deltaPSI (y-axis) versus Branaplam sum deltaPSI (x-axis). Colors and symbols represent sequences found at the -4 or -3 position of the 5’ splice site. There is strong selectivity for -3A (red diamonds) with branaplam, -4A (green x’es) with risdiplam and equipotency for -4-3A (blue squares) with both compounds. B. Same as A, but in cells with U1C knockdown. Comparison between panels A and B demonstrates no change to the sequence selectivity upon U1C siRNA knockdown.

**Supplemental Fig. 8.**
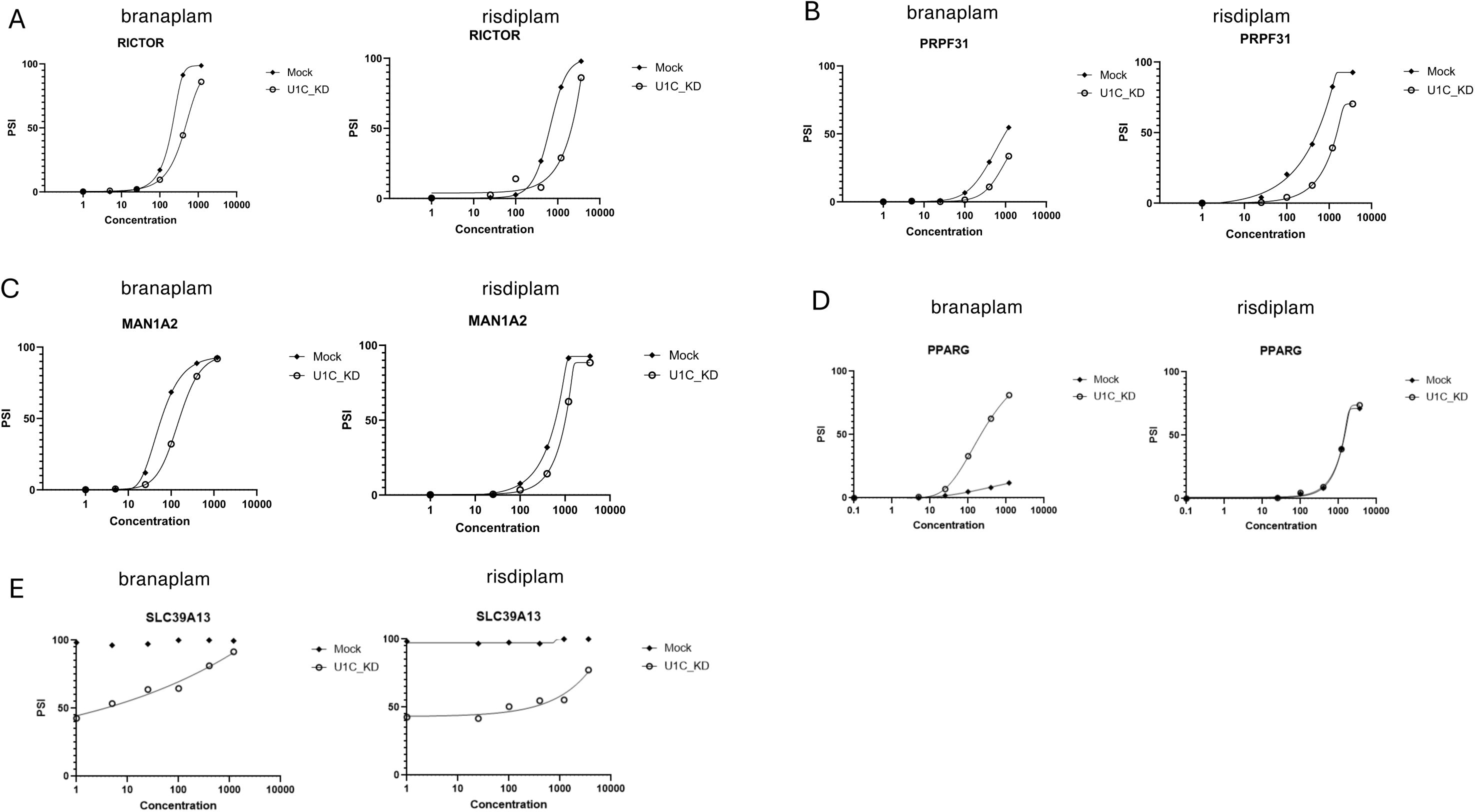
Additional examples demonstrate the effect of U1C KD on iExon inclusion mediated by branaplam or risdiplam. A-C. Plot of PSI of representative exons induced by splicing modulator compound treatment where the inclusion of these exons by compound is reduced in U1C knockdown. All concentrations in Supplemental Fig. 8 are given in nM. D. Plot of PSI of a representative class II exon induced by splicing modulator compound treatment, which represents a very low inclusion exon where splicing inclusion by branaplam is greatly enhance by U1C knockdown. E. Plot of PSI of a representative class III exon induced by splicing modulator compound treatment, where U1C knockdown results in skipping of the exon that is restored when compound is added.

**Supplemental Fig. 9.**
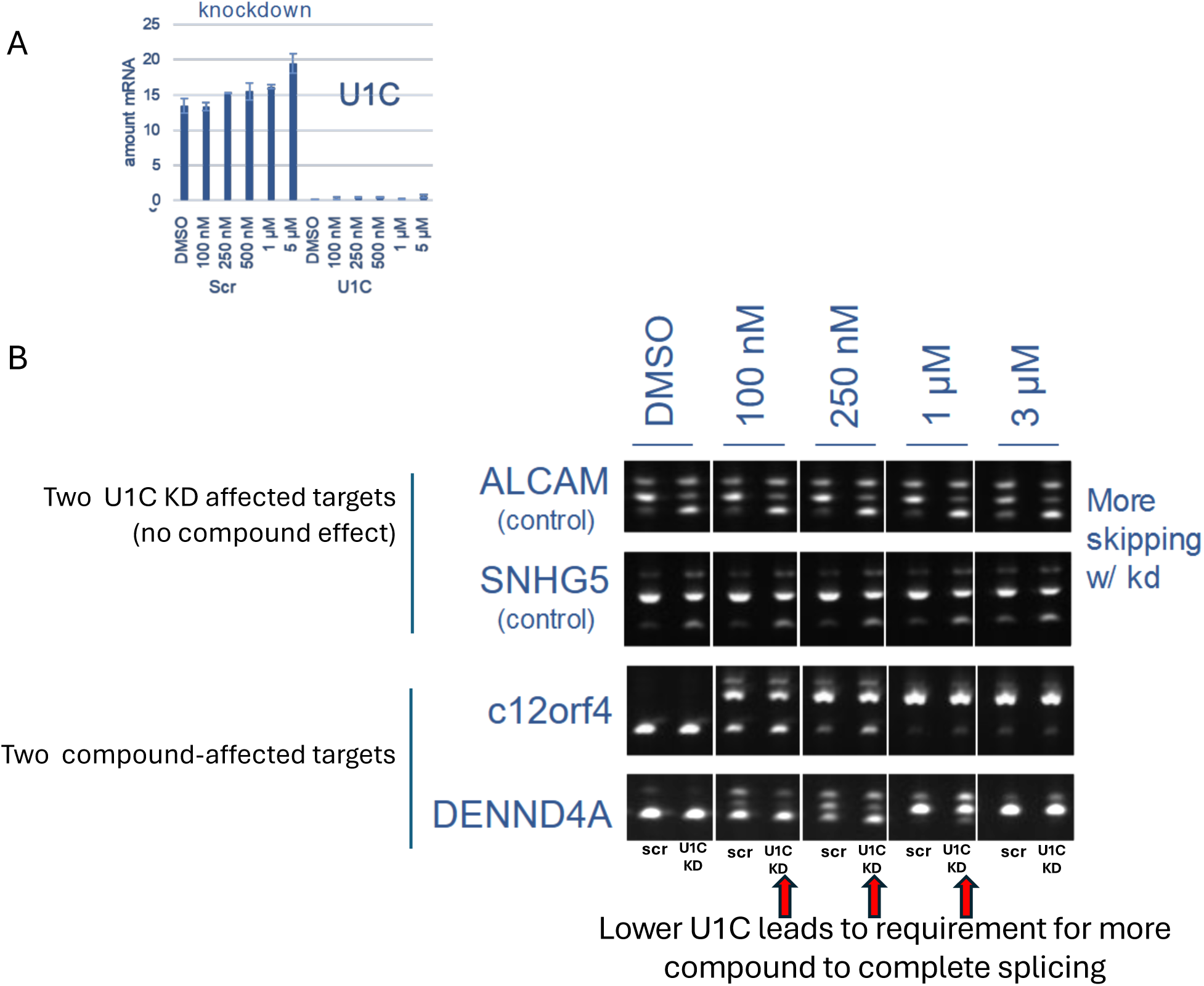
U1C Knockdown reduces branaplam potency as demonstrated by end-point PCR analysis. **A.** qPCR measurement of U1C mRNA after siRNA knockdown with scrambled siRNA control and U1C siRNA. **B.** End-point PCR for known U1C knockdown affected targets ALCAM and SNHG5 and GA/GU targets of branaplam (c12orf4 iExon 1a and DENND4A) scr control versus U1C KD with compound treatment at 3uM, 1uM, 250nm, 100nM and DMSO control. Red arrows depict the doses at which U1C KD effect on branaplam treatment is observed.

**Supplemental Table 1.** Splicing analysis table of RNA-Seq of U1-GA variants with or without U1 helix H mutations. Column names: “coordinates”, the boundary of the exon or exonic region in human genome (hg19); “gene_id”, the NCBI gene ID; “refseqid”, Refseq transcript ID; “reg_len”, length (bp) of the exon or exonic region; “AS_type”, alternative splicing type (exon, cassette exon, the region starts with a 3’ splice sites (ss) and ends with a 5’ss; A5SS, alternative 5’ss region; A3SS, alternative 3’ss region); “startSS_seq” and “endSS_seq”, sequences of start splice site and the end splice site positions (lowercase for intronic sequence and uppercase for exonic sequence). “PSI_SampleName”, Percent-spliced-in (PSI) of an exon or region of a sample or average value of a sample group. “deltaPSI_TestSample_ControlSample”, difference of average PSI values comparing two sample groups (test vs. control); “Pfisher_TestSample_ControlSample”, Fisher’s Exact test *P* value for inclusion and skipping read counts comparing two sample groups (test vs. control); “ReguType_TestSample_ControlSample”, region regulation type. (UP, inclusion; DN, skipping; NC, not statistically significant change; na, not available, eg. lowly expressed genes). HelixH mutations Break1, Break2, ID1, ID2 are G12C, C122G, G12A/C122U, G12C/C122G respectively.

**Supplemental Table 2.** Splicing analysis table of RNA-Seq of risdiplam or branaplam treatment in dose response with U1C knockdown or negative control knockdown in HEK293 cells. See the description for Supplemental Table 1 for column names. P905, branaplam; P683, risdiplam.

## Data availability

The RNA-Seq data was deposited to GEO database under accession GSE304951.

## Acknowledgement

This work was supported by NIH R35GM145289 (R.Z.) and R21NS142950 (R.Z.), and R35GM147498 (J.W.). We would like to thank the Drug Discovery and Development Shared Resource (D^3^SR) for support of the FP experiments. The D^3^SR is supported in part by the CU Cancer Center, an NIH NCI designated cancer center (P30CA046934); and the CU AMC Center for Drug Discovery, which was established from a generous gift from the ALSAM Foundation and through CU AMC institutional support. Molecular graphics were performed with the UCSF Chimera and ChimeraX, developed by the Resource for Biocomputing, Visualization, and Informatics at the University of California, San Francisco, with support from NIGMS P41-GM103311 (Chimera, ChimeraX) and NIH R01-GM129325 (ChimeraX). We thank members of the Zhao lab, especially Dr. Da Cui, for their helpful feedback. We also thank Drs. Frederic Vaillancourt and Bryan Dunyak from Remix Therapeutics for insightful discussion.

