## Supplemental information for "The U1 snRNP protein U1C and Helix H of U1 snRNA are critical for small molecule splicing modulator function"

### Synthesis of branaplam analogs

**General Methods.** Reagents and solvents were purchased from commercial sources (Fisher, Sigma-Aldrich and Combi-Blocks) and used as received. Reactions were tracked by TLC (Silica gel 60 F_254_, Merck) and Agilent 1290 Infinity II HPLC-MS system (Agilent 1290 Infinity II in tandem with LC/MSD iQ Mass Detector). Intermediates and products were purified by a Teledyne ISCO Combi-Flash system using prepacked SiO_2_ cartridges. NMR spectra were acquired on a Bruker AV400 instrument (400 MHz for ^1^H NMR, 100 MHz for ^13^C NMR). ^13^C shifts were obtained with ^1^H decoupling. MestReNova 14.0.1 developed by MESTRELAB RESEARCH was used for NMR data processing. Multiplicity: s, singlet; d, doublet; t, triplet; m, multiplet; brs, broaden singlet. MS-ESI spectra were recorded on Agilent LC/MSD iQ Mass Detector.

Compound **SM2** was synthesized following procedures reported in literature and verified by NMR and Mass spectra^1^.

Synthesis of common intermediate:

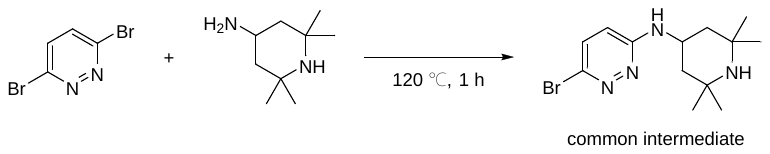

A mixture of 3,6-dibromopyridazine (740 mg, 3.1 mmol) and 2,2,6,6-tetramethylpiperidin-4-amine (970 mg, 6.2 mmol) in a sealed tube was heated at 120 ℃ for 1 h. The resulting crude solid product was purified by silica gel chromatography eluting with 0-30% methanol in dichloromethane to afford the pure product as light yellow solid (500 mg, yield 51.5%). MS-ESI (*m*/*z*) [M+H]^+^ 313.1.

General procedure for synthesizing the final product:

A mixture of the common intermediate (30 mg, 0.095 mmol), aryl borate ester (2 eq, 0.19 mmol), Pd(PPh_3_)_4_ (0.1 eq, 0.0095 mmol), K_2_CO_3_ (3 eq, 0.285 mmol) and dioxane/water (1.5 mL/0.3 mL) in a microwave tube was degassed and refilled with nitrogen. The mixture was then heated at 100 ℃ for 2 h in a Biotage initiator+ microwave reactor. The resulting reaction mixture was filtered and concentrated, the crude product was purified by silica gel chromatography eluting with 0-30% methanol in dichloromethane to afford the pure product.

6-phenyl-N-(2,2,6,6-tetramethylpiperidin-4-yl)pyridazin-3-amine (**A9**)

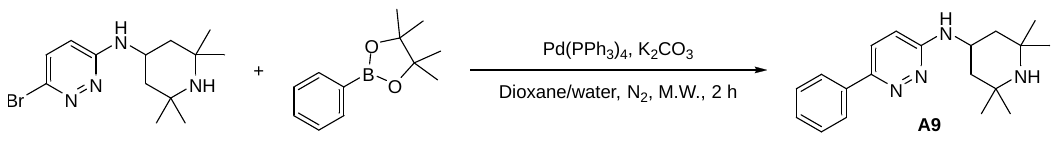

Following the General procedure for synthesizing the final product, 20 mg white solid was obtained (yield 67.2%). MS-ESI (*m*/*z*) [M+H]^+^ 311.2.

^1^H NMR (400 MHz, DMSO-*d*_6_) δ 8.75 (s, 1H), 7.98 (d, *J* = 7.1 Hz, 2H), 7.85 (d, *J* = 9.4 Hz, 1H), 7.48 (t, *J* = 7.5 Hz, 2H), 7.40 (t, *J* = 7.2 Hz, 1H), 7.09 (d, *J* = 7.4 Hz, 1H), 6.93 (d, *J* = 9.3 Hz, 1H), 4.57 – 4.43 (m, 1H), 2.16 (dd, *J* = 13.6, 3.6 Hz, 2H), 1.51 – 1.47 (m, 8H), 1.44 (s, 6H).

^13^C NMR (100 MHz, DMSO-*d*_6_) δ 157.7, 150.6, 137.4, 129.2, 128.9, 125.8, 125.6, 115.8, 74.0, 57.4, 41.5, 41.2, 30.4, 25.4, 24.9.

3-(6-((2,2,6,6-tetramethylpiperidin-4-yl)amino)pyridazin-3-yl)naphthalen-2-ol (**A10**)

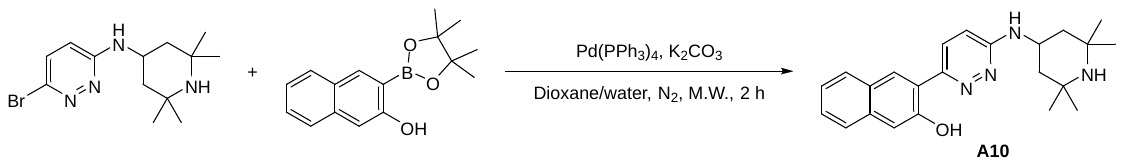

Following the General procedure for synthesizing the final product, 10 mg light yellow solid was obtained (yield 27.7%). MS-ESI (*m*/*z*) [M+H]^+^ 377.2.

^1^H NMR (400 MHz, DMSO-*d*_6_) δ 13.59 (s, 1H), 8.45 (s, 1H), 8.28 (d, *J* = 9.7 Hz, 1H), 7.87 (d, *J* = 8.2 Hz, 1H), 7.71 (d, *J* = 8.3 Hz, 1H), 7.44 (ddd, *J* = 8.2, 6.7, 1.3 Hz, 1H), 7.33 – 7.28 (m, 2H), 7.14 (d, *J* = 7.7 Hz, 1H), 7.09 (d, *J* = 9.6 Hz, 1H), 4.43 – 4.35 (m, 1H), 1.93 (dd, *J* = 12.4, 3.7 Hz, 2H), 1.25 (s, 6H), 1.11 – 1.05 (m, 8H).

^13^C NMR (100 MHz, DMSO-*d*_6_) δ 157.6, 156.0, 151.7, 134.9, 128.6, 127.7, 127.4, 126.5, 126.3, 126.0, 123.6, 121.2, 118.0, 111.3, 51.3, 44.6, 43.7, 34.9, 29.0.

6-(quinolin-3-yl)-N-(2,2,6,6-tetramethylpiperidin-4-yl)pyridazin-3-amine (**A13**)

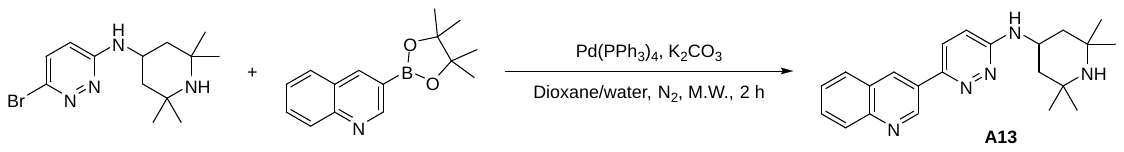

Following the General procedure for synthesizing the final product, 14.5 mg light yellow solid was obtained (yield 41.9%). MS-ESI (*m*/*z*) [M+H]^+^ 362.2.

^1^H NMR (400 MHz, DMSO-*d*_6_) δ 9.61 (d, *J* = 2.3 Hz, 1H), 8.88 (d, *J* = 1.5 Hz, 1H), 8.10 – 8.06 (m, 3H), 7.95 (brs, 1H), 7.79 (ddd, *J* = 8.5, 6.9, 1.5 Hz, 1H), 7.67 (ddd, *J* = 8.1, 6.9, 1.2 Hz, 1H), 7.26 (d, *J* = 7.3 Hz, 1H), 7.02 (d, *J* = 9.4 Hz, 1H), 4.57 – 4.53 (m, 1H), 2.18 (dd, *J* = 13.6, 3.6 Hz, 2H), 1.57 – 1.51 (m, 8H), 1.45 (s, 6H).

^13^C NMR (100 MHz, DMSO-*d*_6_) δ 157.9, 148.7, 148.6, 147.7, 132.1, 130.3, 130.1, 129.2, 129.0, 128.0, 127.6, 126.0, 115.9, 57.3, 41.6, 41.1, 30.4, 24.9.

6-(isoquinolin-6-yl)-N-(2,2,6,6-tetramethylpiperidin-4-yl)pyridazin-3-amine (**A14**)

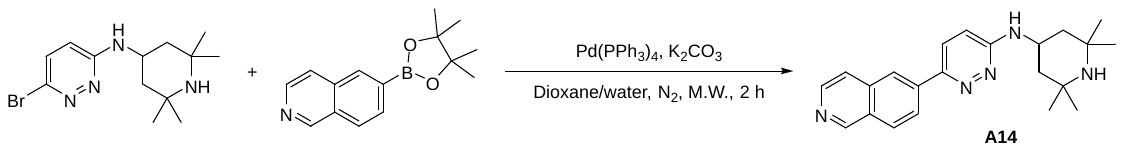

Following the General procedure for synthesizing the final product, 22 mg light yellow solid was obtained (yield 63.5%). MS-ESI (*m*/*z*) [M+H]^+^ 362.2.

^1^H NMR (400 MHz, DMSO-*d*_6_) δ 9.34 (s, 1H), 8.76 (s, 1H), 8.55 – 8.53 (m, 2H), 8.41 (dd, *J* = 8.6, 1.7 Hz, 1H), 8.21 (d, *J* = 8.7 Hz, 1H), 8.08 (d, *J* = 9.4 Hz, 1H), 7.89 (d, *J* = 5.8 Hz, 1H), 7.27 (d, *J* = 7.3 Hz, 1H), 7.00 (d, *J* = 9.4 Hz, 1H), 4.56 – 4.54 (m, 1H), 2.18 (dd, *J* = 13.6, 3.6 Hz, 2H), 1.56 – 1.50 (m, 8H), 1.44 (s, 6H).

^13^C NMR (100 MHz, DMSO-*d*_6_) δ 158.0, 152.6, 149.7, 143.9, 139.0, 136.0, 128.6, 128.3, 126.1, 125.4, 123.0, 121.2, 115.7, 57.4, 41.6, 41.1, 30.4, 24.9.

NMR Spectra

Compound A9

^1^H NMR

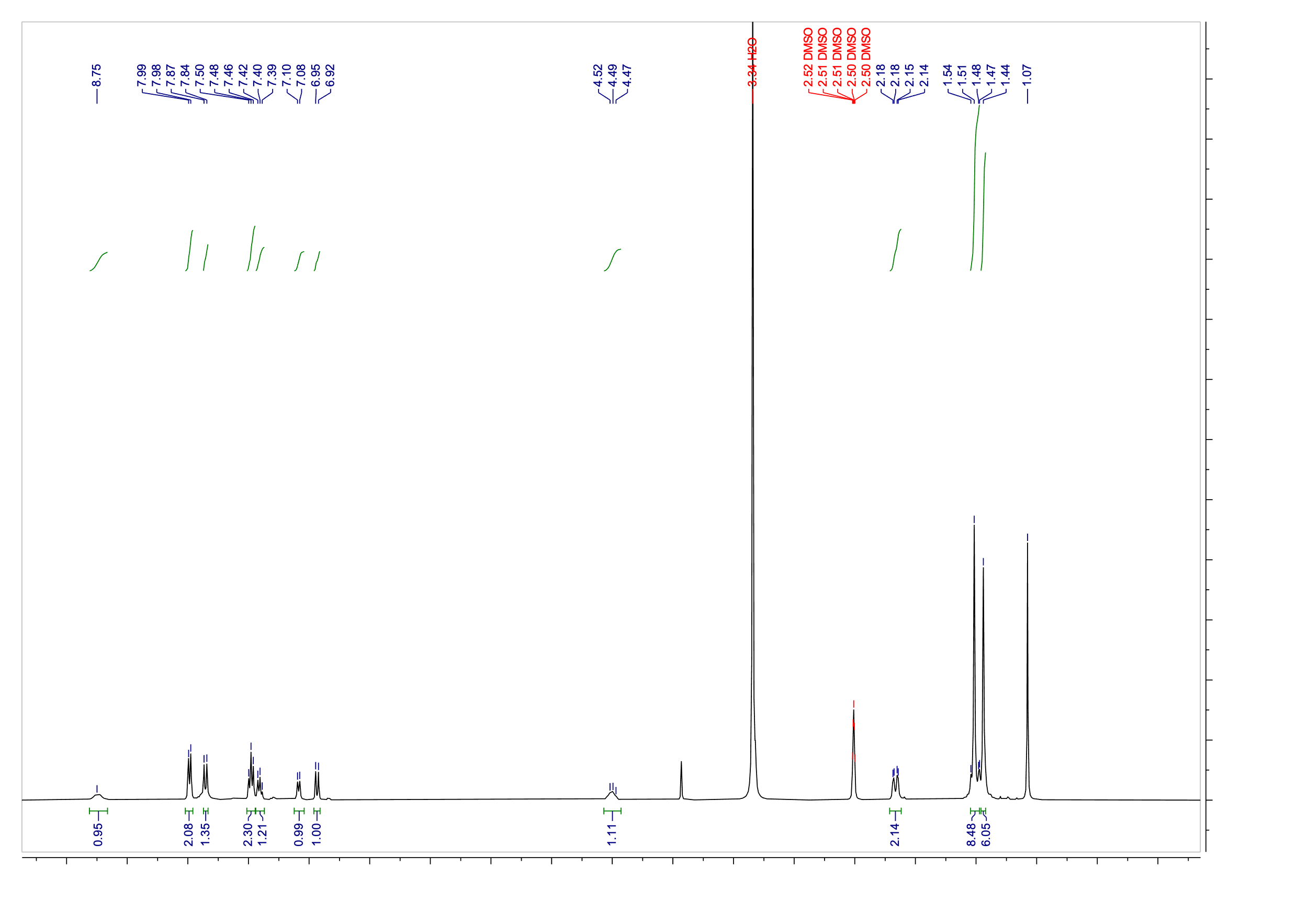

^13^C NMR

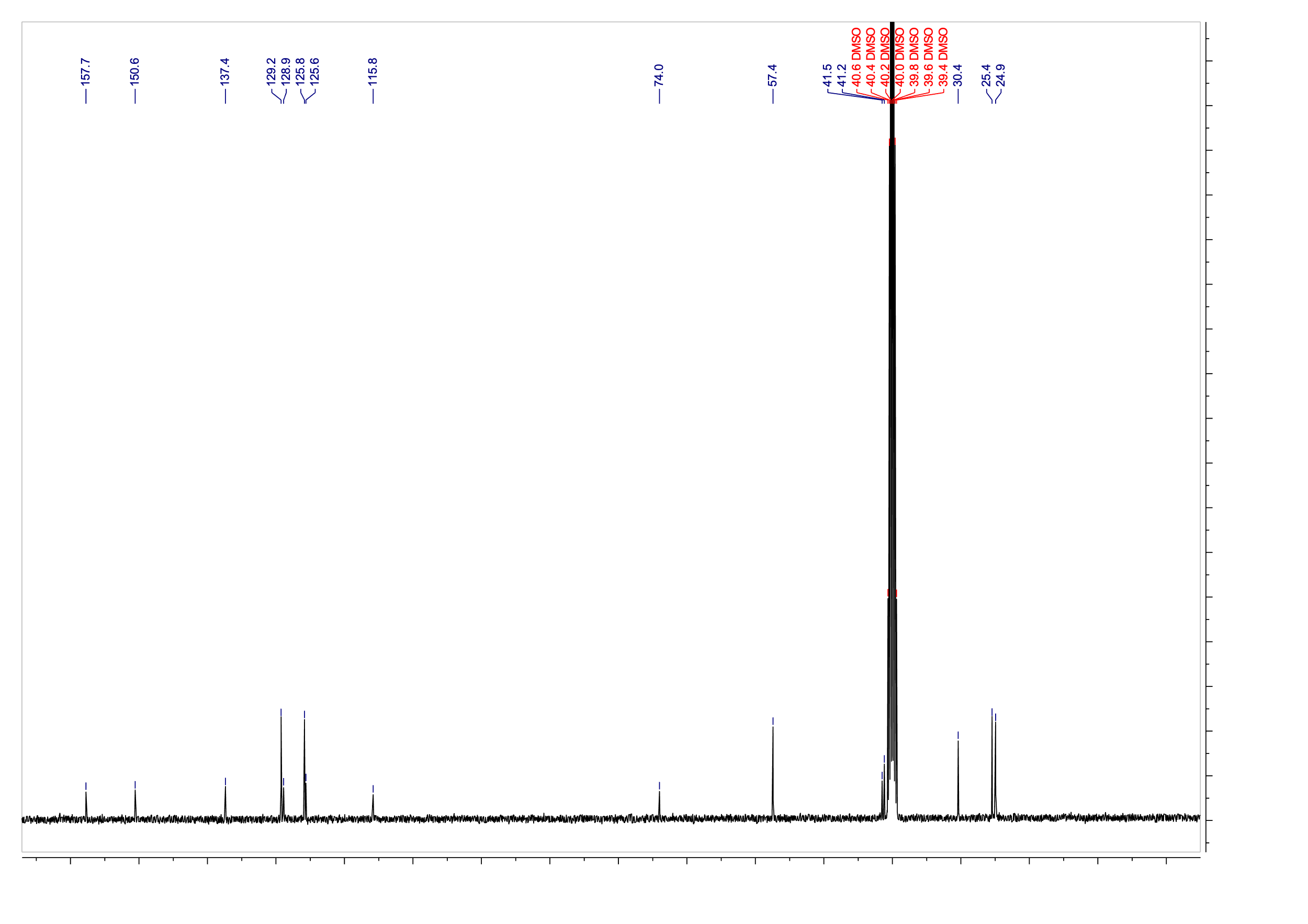

Compound A10

^1^H NMR

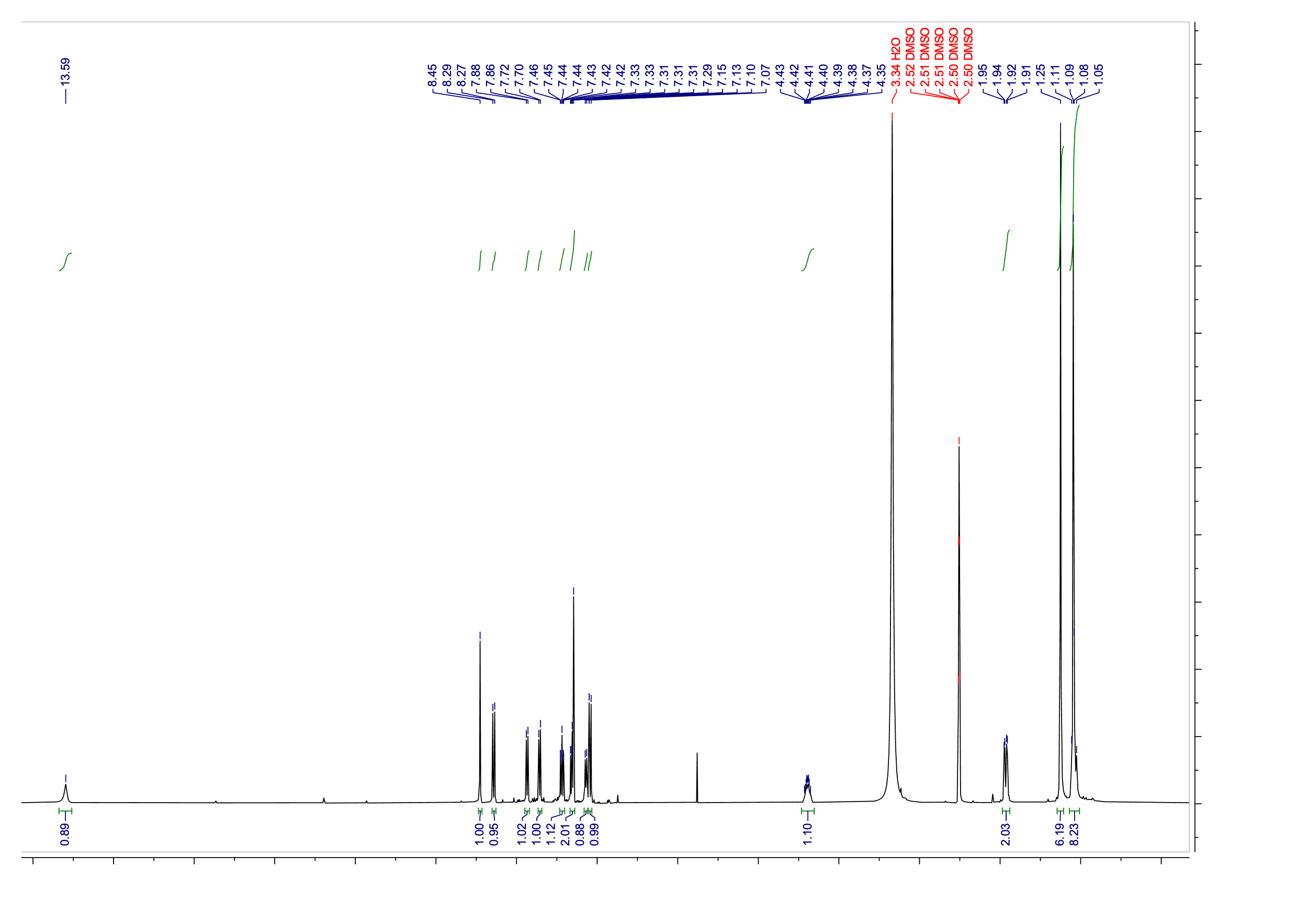

^13^C NMR

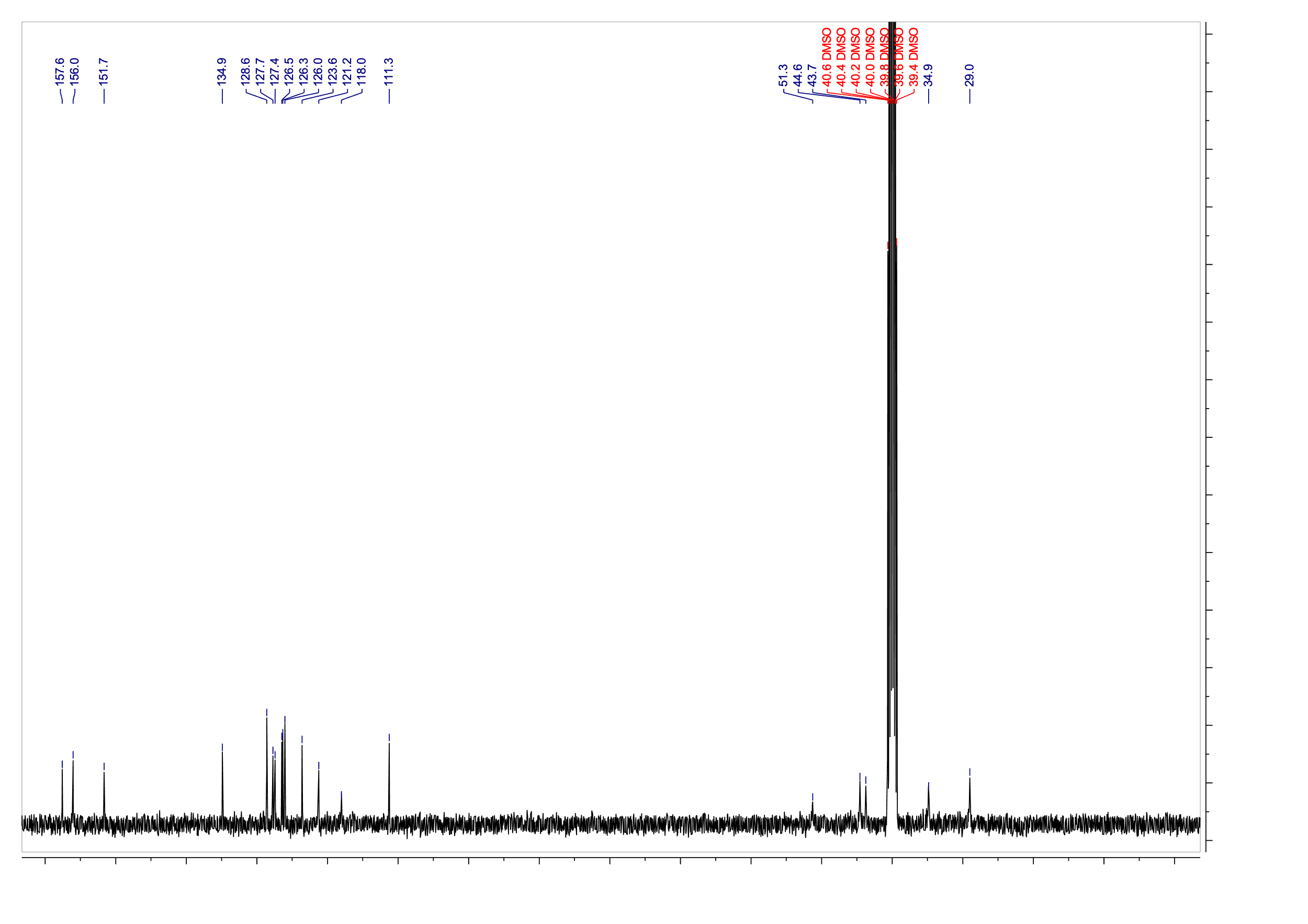

Compound A13

^1^H NMR

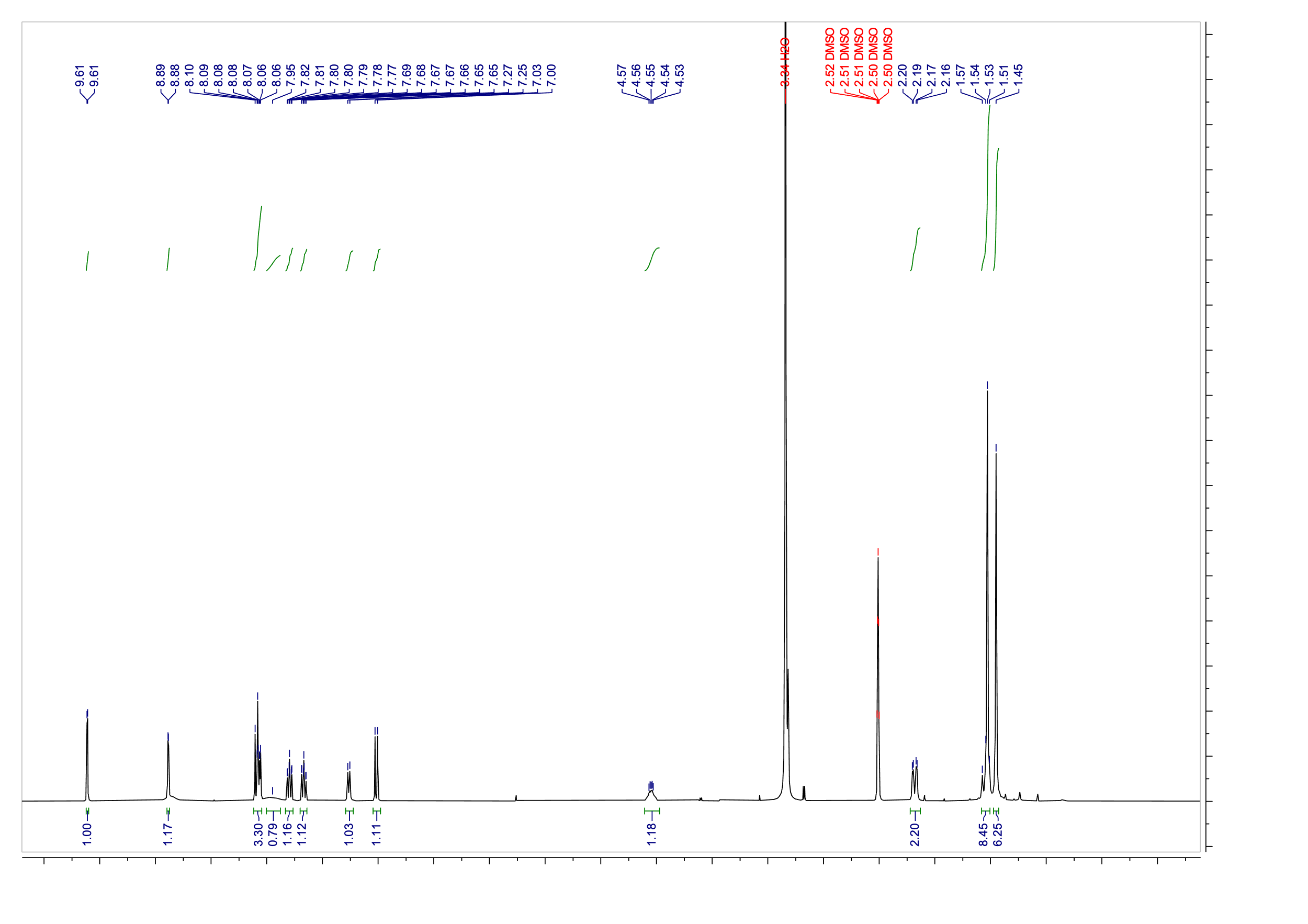

^13^C NMR

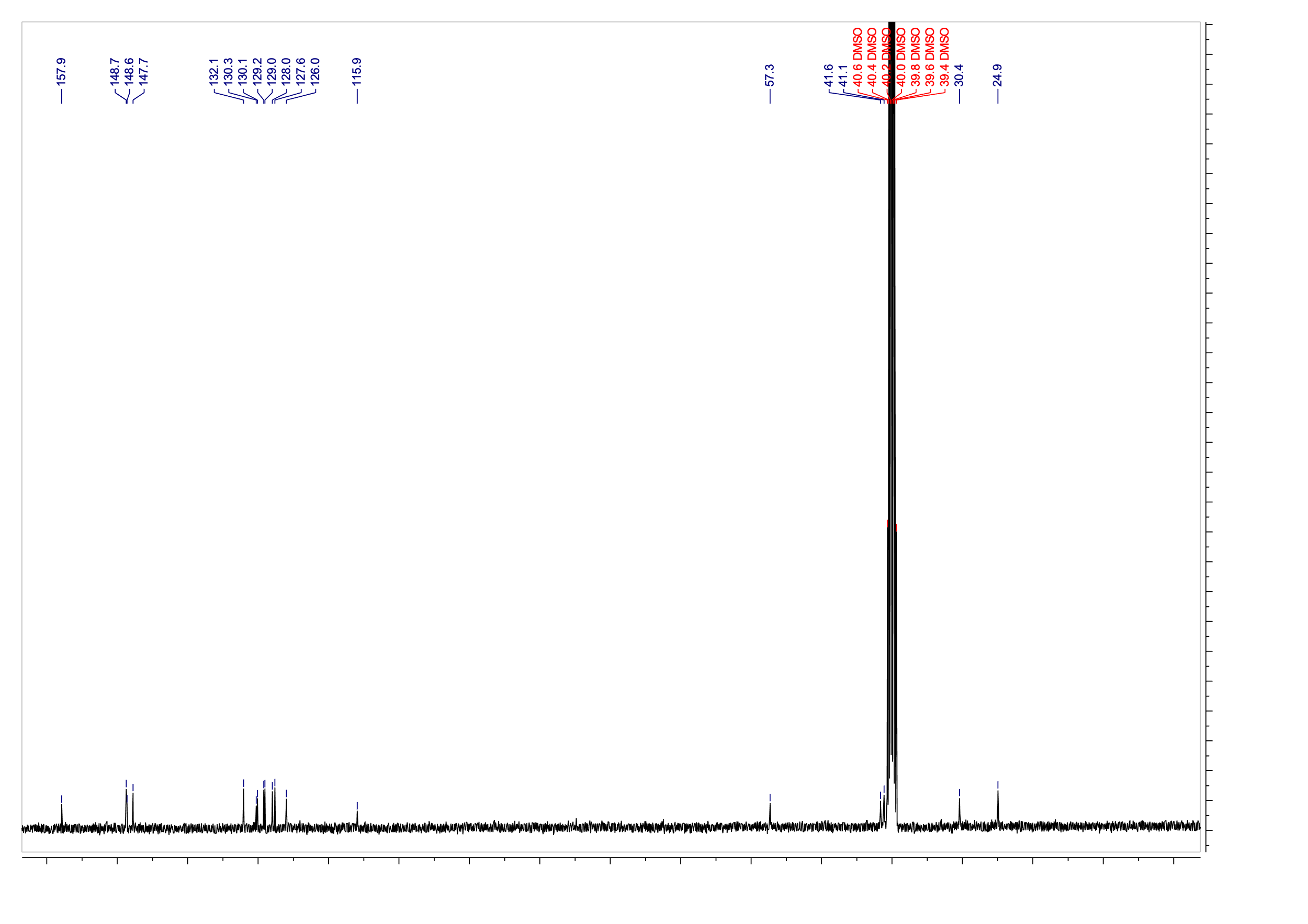

Compound A14

^1^H NMR

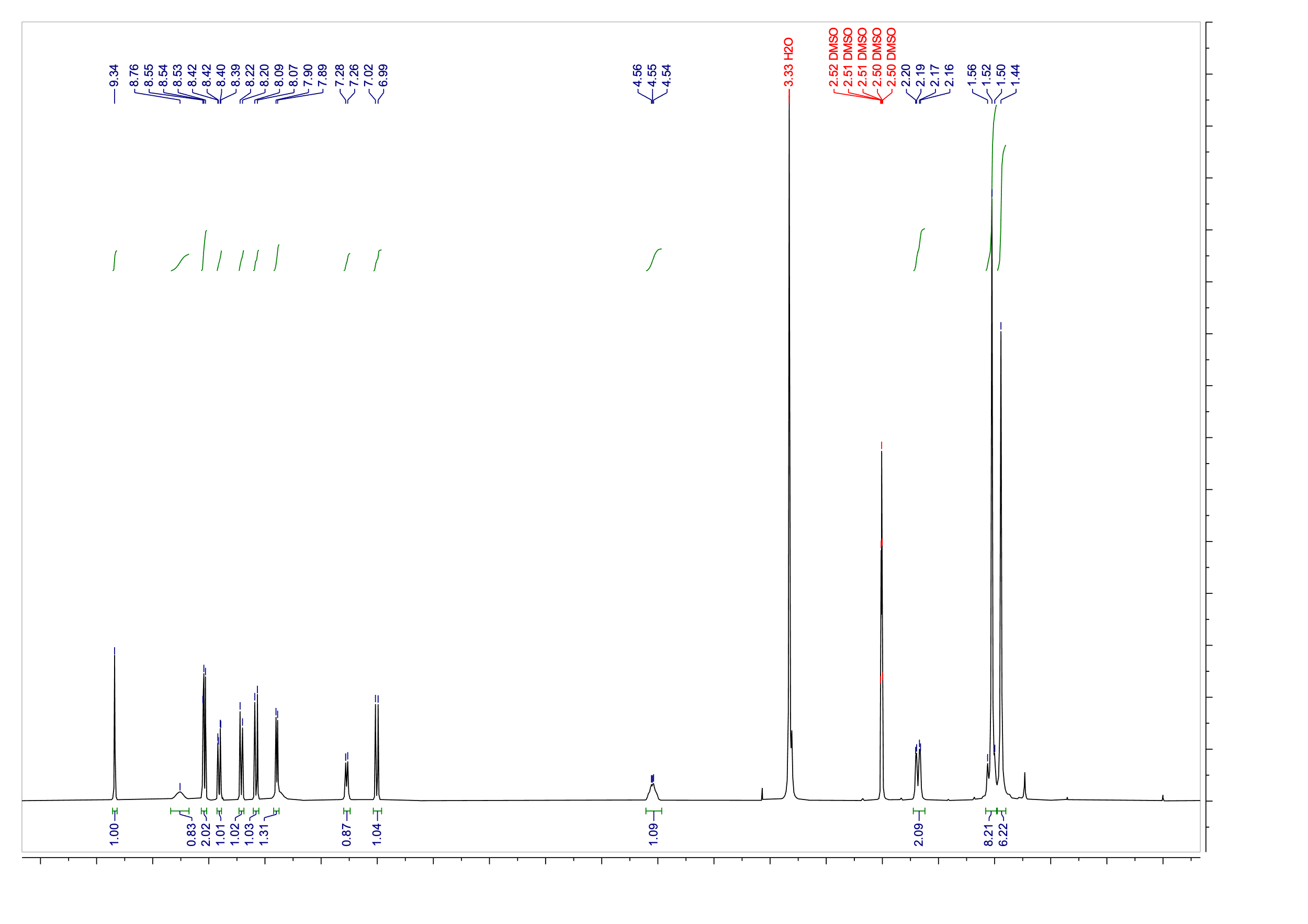

^13^C NMR

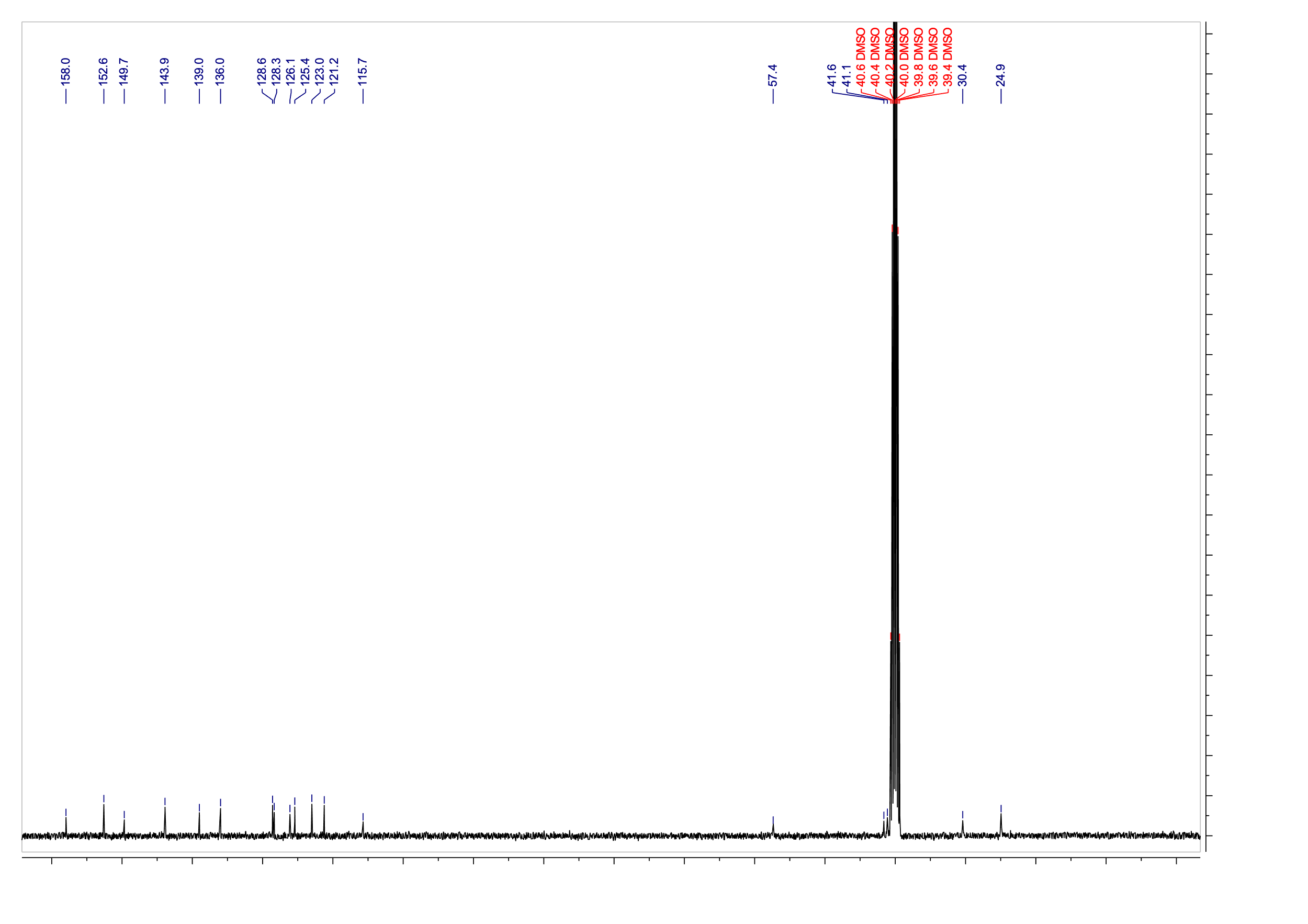
